## Supplementary Information for "A synthetic tubular molecular transport system"

[2]Lehrstuhl für Theoretische Bio- und Softmatter Physik, Physik Department, Freie Universität Berlin, Berlin, Germany

#### Supplementary Information

##### Contents:

Supplementary Figures 1-41
Supplementary Note 1
Materials & Methods
Captions to Supplementary Movies

### Supplementary Figures

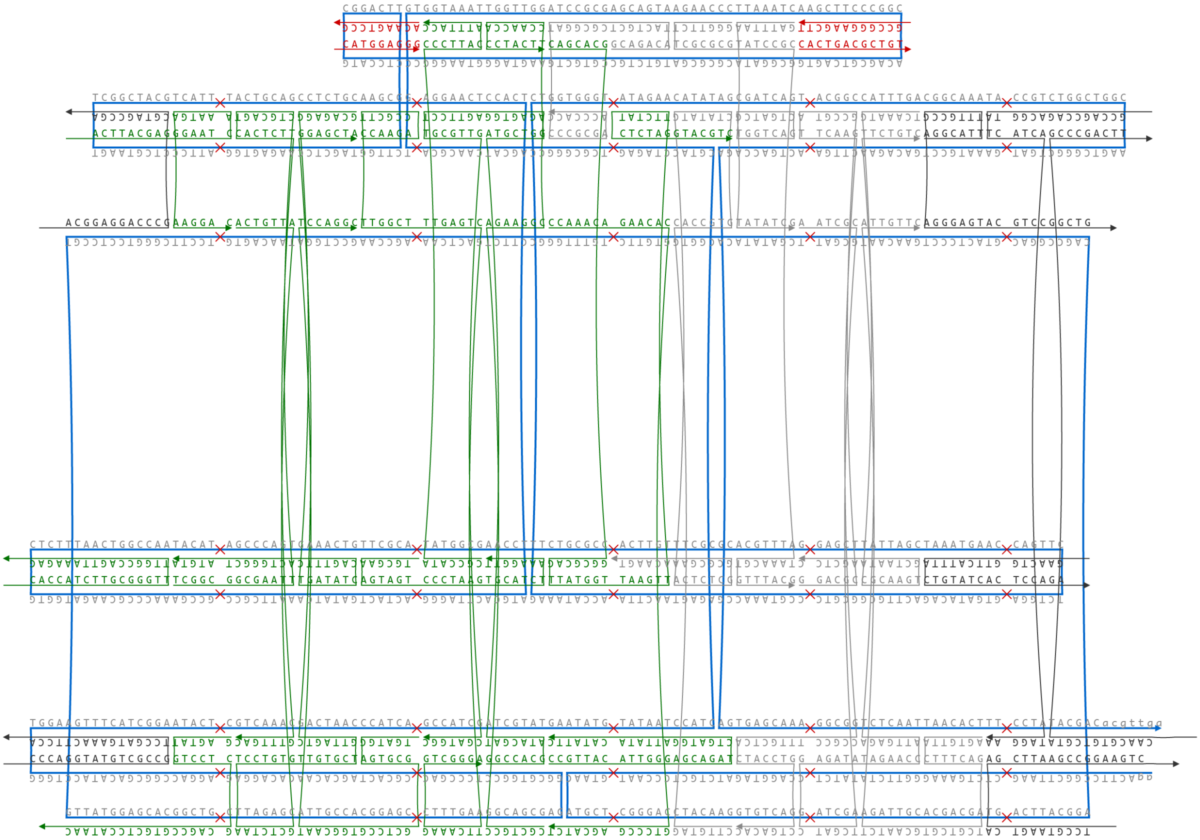

**Figure S 1** Strand diagram of the piston. Made with caDNAno v0.2 (*1*).

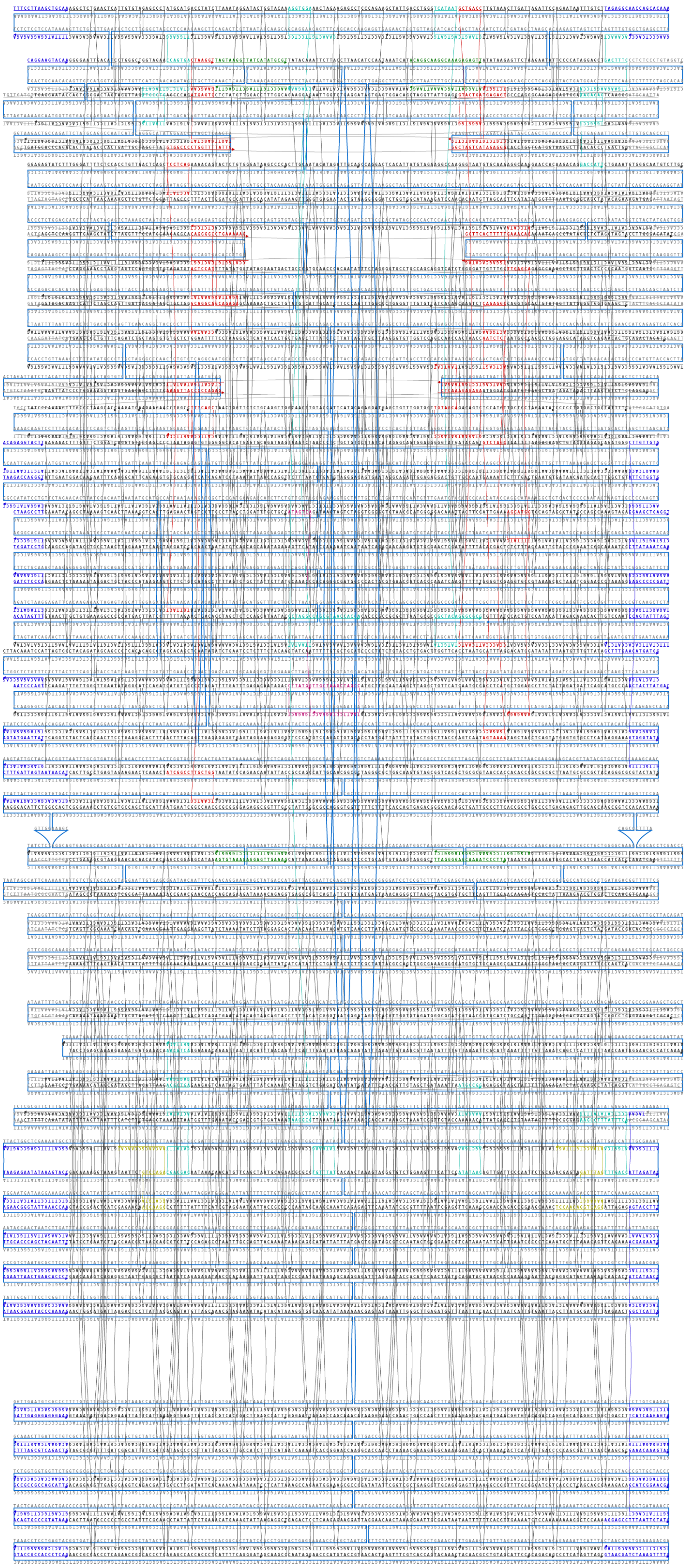

**Figure S 2** Strand diagram of the barrel. Made with caDNAno v0.2 (*1*).

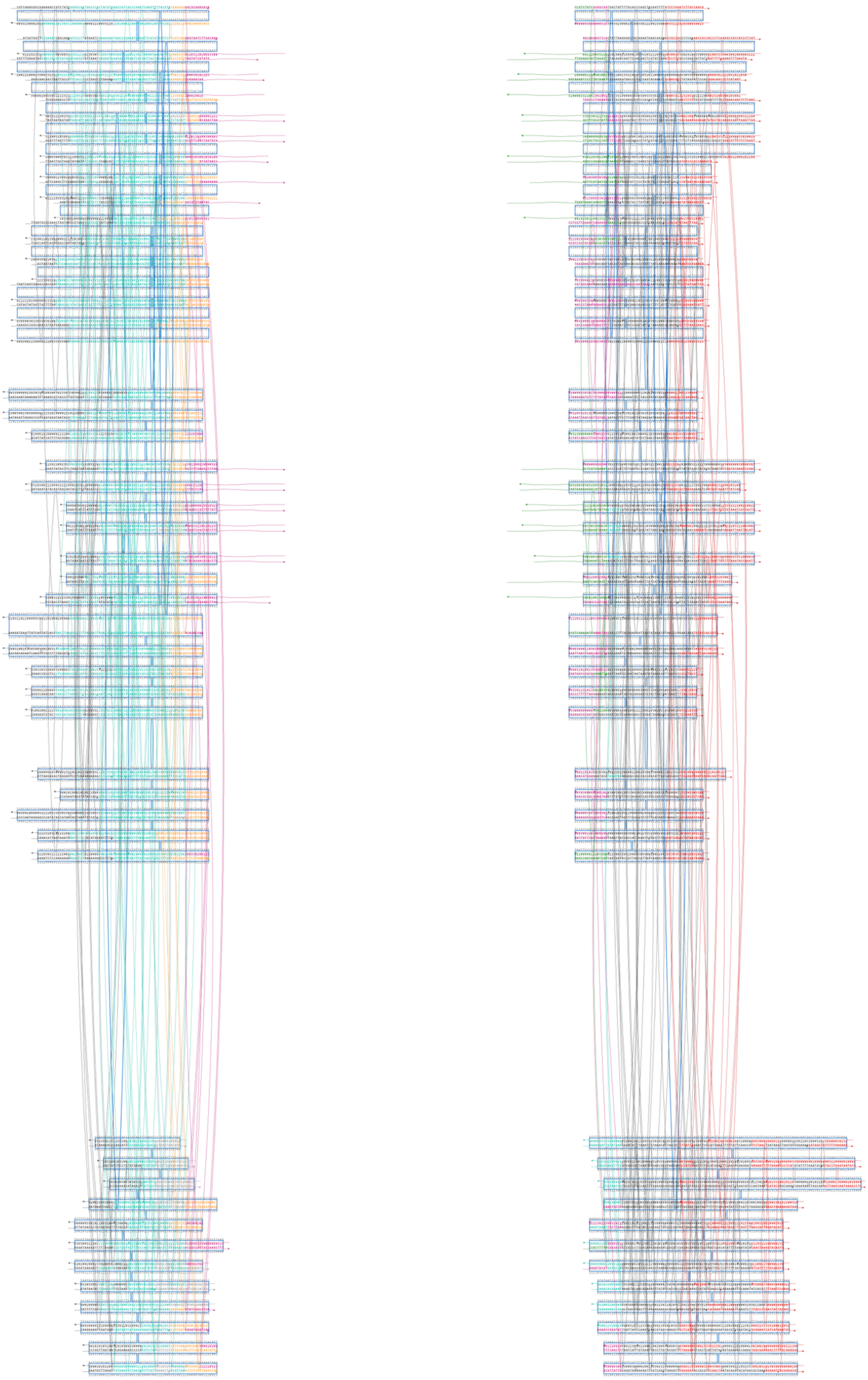

**Figure S 3** Strand diagrams of the cap building blocks (left: cap 1, right: cap 2). Made with caDNAno v0.2 (*1*).

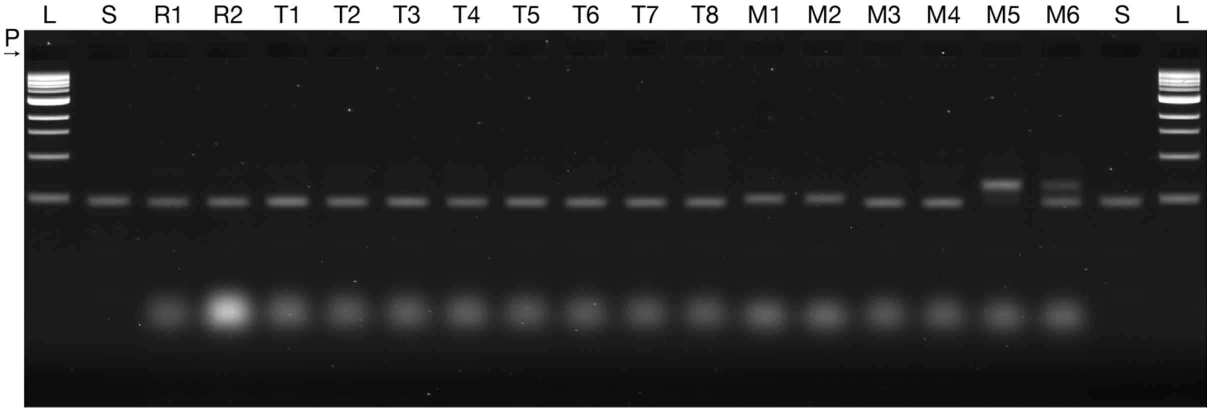


**Figure S 4 Piston folding screen.** Laser-scanned image of a 2.5% agarose gel with 5.5 mM $MgCl_2$ run on a water bath at 90 V for 90 min on which the following samples were electrophoresed: L, 1kb ladder; S, 1033 bases long scaffold; R1, folding interval 60-44°C, 20 mM $MgCl_2$, 5x staple-to-scaffold excess; R2, 60-44°C, 20 mM $MgCl_2$, 10x staple-to-scaffold excess; lanes T1-T8: 20 mM $MgCl_2$; T1, folding interval 50-47°C; T2, folding interval 52-49°C; T3, folding interval 54-51°C; T4, folding interval 56-53°C; T5, folding interval 58-55°C; T6, folding interval 60-57°C; T7, folding interval 62-59°C; T8, folding interval 64-61°C; lanes M1-M6, folding interval 60-44°; M1, 5 mM $MgCl_2$; M2, 10 mM $MgCl_2$; M3, 15 mM $MgCl_2$; M4, 20 mM $MgCl_2$; M5, 25 mM $MgCl_2$; M6, 30 mM $MgCl_2$. P indicates the pockets.

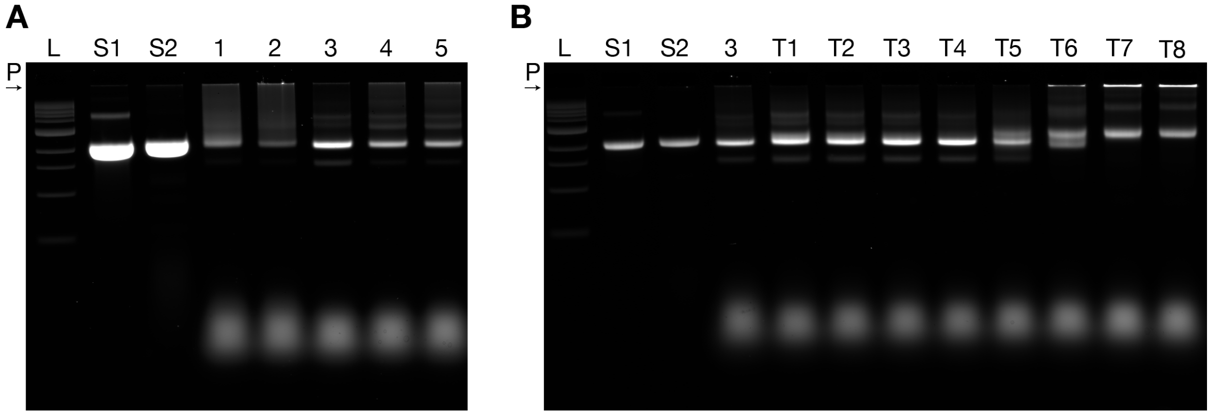


**Figure S 5 Barrel folding screens.** Laser-scanned image of 2 **(A and B)** 1.5% agarose gels with 5.5 mM $MgCl_2$ run on a water bath at 90 V for 90 min on which the following samples were electrophoresed: L, 1kb ladder; S1, 7560 bases long scaffold type 1; S2, 7560 bases long scaffold type 2 (orthogonal to S1); 1-5 and T1-T8, barrel folding reactions, all starting with 15 min at 65°C; 1, 20 mM $MgCl_2$, 60-44°C 1 hour/°C; 2, 20 mM $MgCl_2$, 60-44°C 2 hours /°C; 3, 15 mM $MgCl_2$, 60-44°C 3 hours/°C; 4, 20 mM $MgCl_2$, 60-44°C 3 hours/°C; 5, 25 mM $MgCl_2$, 60-44°C 3 hours/°C; T1-T8: 15 mM $MgCl_2$, 3 hours/°C; T1, 50-47°C; T2, 52-49°C; T3, 54-51°C; T4, 56-53°C; T5, 58-55°C; T6, 60-57°C; T7, 62-59°C; T8, 64-61°C. P indicates the pockets.

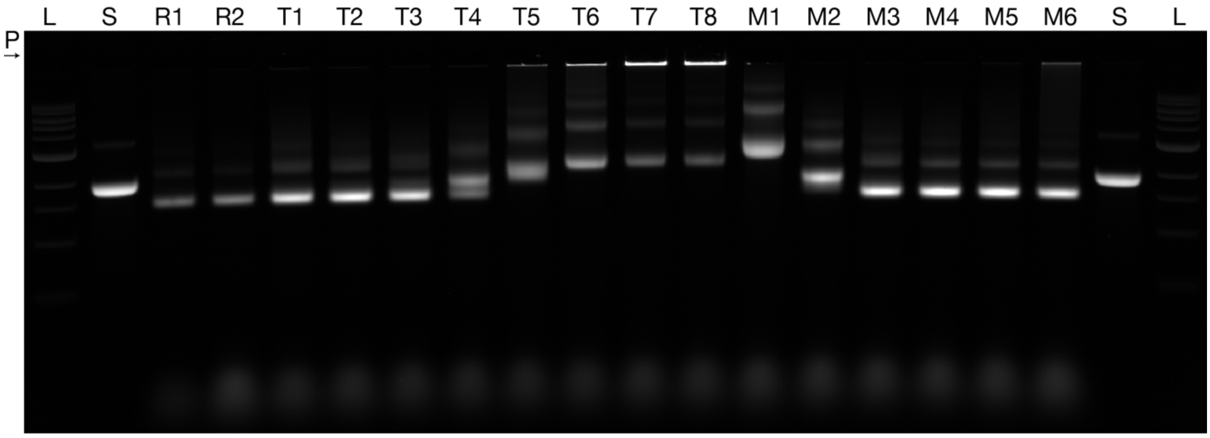


**Figure S 6 Cap1 folding screen.** Laser-scanned image of a 2% agarose gel with 5.5 mM $MgCl_2$ run on a water bath at 90 V for 90 min on which the following samples were electrophoresed: L, 1kb ladder; S, 7560 bases long scaffold; R1, folding interval 60-44°C, 20 mM $MgCl_2$, 5x staple-to-scaffold excess; R2, 60-44°C, 20 mM $MgCl_2$, 10x staple-to-scaffold excess; lanes T1-T8: 20 mM $MgCl_2$; T1, folding interval 50-47°C; T2, folding interval 52-49°C; T3, folding interval 54-51°C; T4, folding interval 56-53°C; T5, folding interval 58-55°C; T6, folding interval 60-57°C; T7, folding interval 62-59°C; T8, folding interval 64-61°C; lanes M1-M6, folding interval 60-44°; M1, 5 mM $MgCl_2$; M2, 10 mM $MgCl_2$; M3, 15 mM $MgCl_2$; M4, 20 mM $MgCl_2$; M5, 25 mM $MgCl_2$; M6, 30 mM $MgCl_2$. P indicates the pockets.

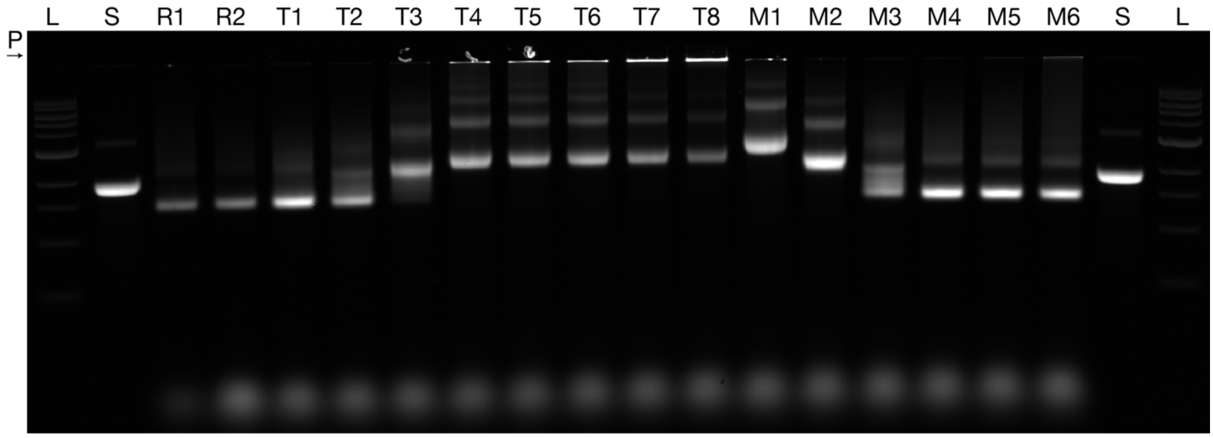


**Figure S 7 Cap2 folding screen.** Laser-scanned image of a 2% agarose gel with 5.5 mM $MgCl_2$ run on a water bath at 90 V for 90 min on which the following samples were electrophoresed: L, 1kb ladder; S, 7560 bases long scaffold; R1, folding interval 60-44°C, 20 mM $MgCl_2$, 5x staple-to-scaffold excess; R2, 60-44°C, 20 mM $MgCl_2$, 10x staple-to-scaffold excess; lanes T1-T8: 20 mM $MgCl_2$; T1, folding interval 50-47°C; T2, folding interval 52-49°C; T3, folding interval 54-51°C; T4, folding interval 56-53°C; T5, folding interval 58-55°C; T6, folding interval 60-57°C; T7, folding interval 62-59°C; T8, folding interval 64-61°C; lanes M1-M6, folding interval 60-44°; M1, 5 mM $MgCl_2$; M2, 10 mM $MgCl_2$; M3, 15 mM $MgCl_2$; M4, 20 mM $MgCl_2$; M5, 25 mM $MgCl_2$; M6, 30 mM $MgCl_2$. P indicates the pockets.

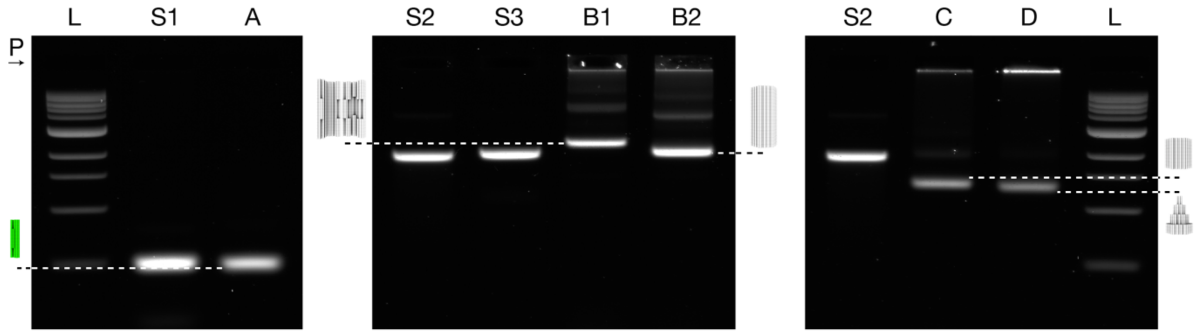


**Figure S 8 EMA quality control of monomers.** Laser-scanned image (split into 3 images) of a 2% agarose gel with 5.5 mM $MgCl_2$ run on a water bath at 90 V for 90 min on which the following samples were electrophoresed: L, 1kb ladder; S1, 1033 bases long scaffold; A, folded and purified piston; S2, 7560 bases long scaffold type 1; S3, 7560 bases long scaffold type 2 (orthogonal to S1); B1, folded and purified barrel in a dynamic/open configuration; B2, folded and purified barrel in a permanently closed configuration; C, folded and purified cap 1; D, folded and purified cap 2. Dashed lines point from the corresponding bands to models of the folded and purified structures. P indicates the pockets.

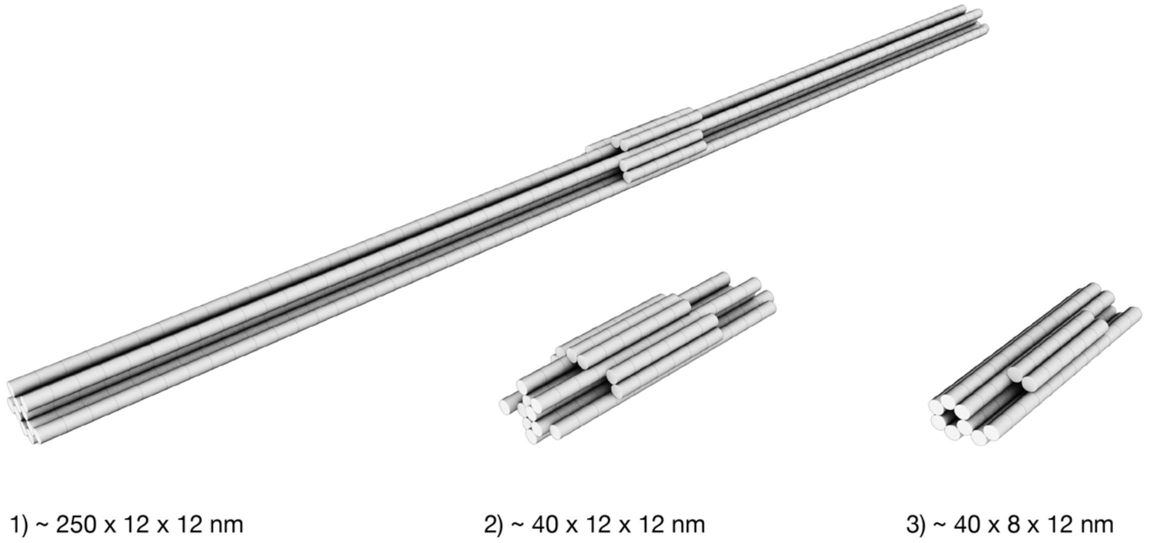


**Figure S 9 Different piston variants.** Models of piston variants. **Left:** variant 1 (250 x 12 x 12 nm). **Middle:** variant 2 (40 x 12 x 12 nm). **Right:** variant 3 (40 x 8 x 12 nm). Cylinders represent DNA double-helices.

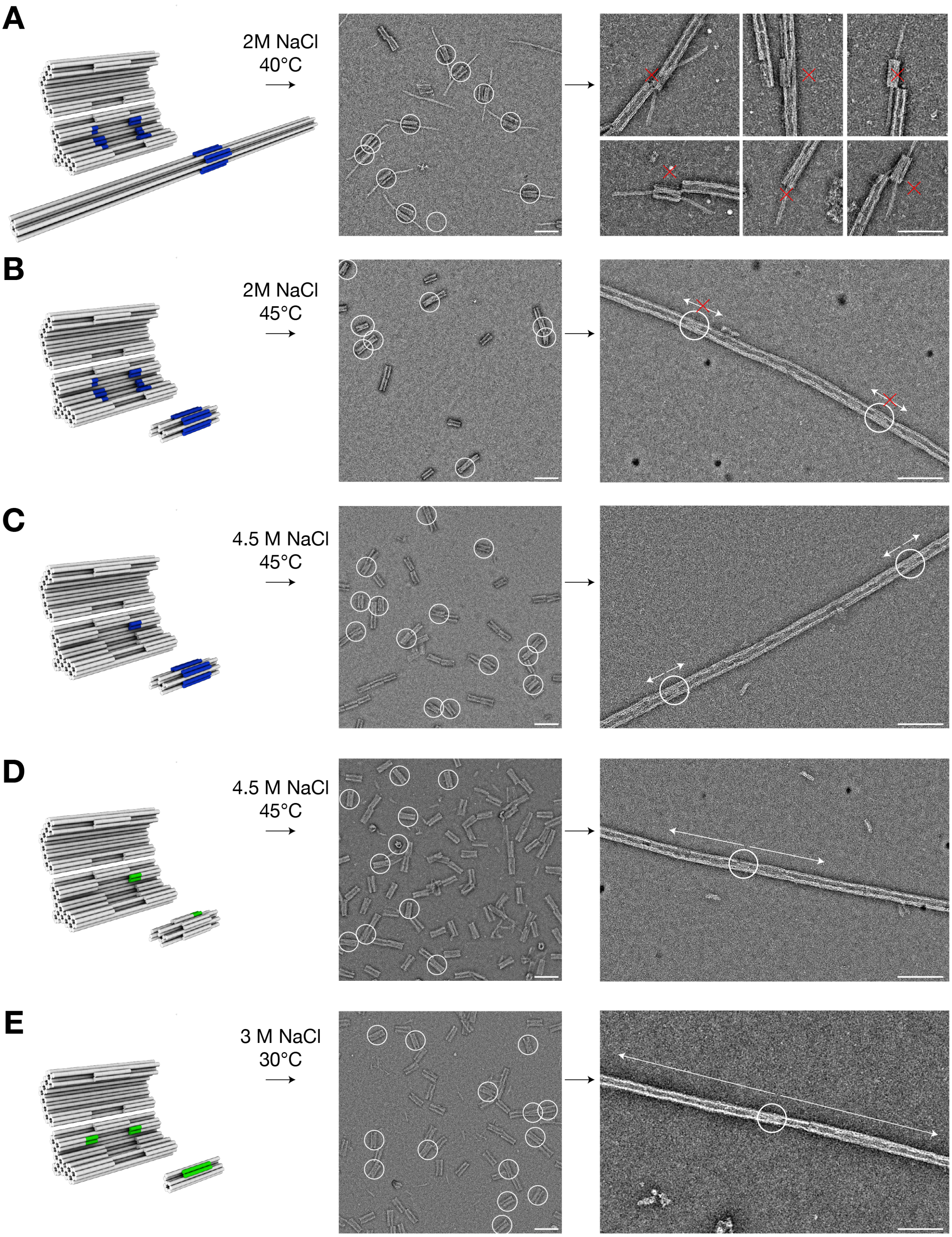


**Figure S 10 Loading the tunnels with different piston variants. (A-E) Left:** models of the barrel and piston variants. Cylinders represent DNA double-helices. The barrel is illustrated in an open conformation. Blue and green protrusions and recesses on the barrel and pistons highlight interaction sites. Blue: blunt-end stacking interactions; green: hybridization of single stranded overhangs and single stranded scaffold loops on the piston and the barrel binding site. **Middle:** typical field of view

TEM micrographs of assembled barrel-piston dimers imaged with a Philips CM100 microscope. White circles highlight correctly assembled dimers. **Right:** typical field of view TEM micrographs of extended tracks containing loaded barrels imaged with a Philips CM100 and an FEI Tecnai120 microscope. White circles highlight pistons inside correctly formed filaments. Red crosses highlight assembly defects that inhibit piston movement. **(A)** Piston variant 1, folded from a 7560 bases long scaffold, is approximately 4 times longer than the barrel in the helical direction. It binds to the barrel binding site by blunt-end stacking interactions. This reaction occurs at 40°C and 2 M NaCl (left). Negative-staining TEM confirms correctly assembled barrel-piston dimers (middle). The filament assembly step leads to defects with the piston often sticking out laterally at barrel-barrel interfaces. We therefore tested a second piston variant **(B)** which is shorter than the barrel and was folded from a 1545 bases long scaffold. Piston variant B was docked to the barrel via blunt-end stacking interactions. This reaction occurs at 45°C and 2 M NaCl (left). Negative-staining TEM confirms correctly assembled barrel-piston dimers (middle). The filament assembly now results in correctly enclosed pistons inside of filaments (right). However, even at very low ionic strengths (0.3-0.5 M NaCl) the piston would not move, which we attributed to failure to undock from the binding site. For variant **(C)**, we reduced the number of blunt-end stacking bonds from 12 down to 2. The other blunt ends were passivated with 5 T overhangs. The dimerization reaction now occurs at 45°C and 4.5 M NaCl (left). Negative-staining TEM confirms correctly assembled barrel-piston dimers (middle). The filament assembly shows correctly enclosed pistons inside of filaments (right). However, real-time fluorescence mobility experiments at low ionic strengths (0.3-0.5 M NaCl) again showed very low mobility of this piston variant. Further reducing the NaCl concentration led to filament disassembly. Reducing the blunt-end stacking interaction between the barrel and piston even further was not practical, since the loading reaction ceased to work. For variant **(D)**, the docking site of the piston now features single stranded scaffold loops whose sequences are complementary to single stranded overhangs in the barrel binding site. The dimerization reaction occurs at 45°C and 4.5 M NaCl (left). Negative-staining TEM confirms correctly assembled barrel-piston dimers (middle). The filament assembly again resulted in correctly enclosed pistons inside of filaments (right). Mobility experiments were performed upon adding invader strands that bind to the piston, thereby releasing it from its initial binding site by toehold-mediated strand displacement. This piston variant showed improved mobility compared to the previous variant (C). However, it still only covered total displacements of up to several 100 nm. For variant **(E)**, we reduced the cross section of the piston to make its motion less susceptible to potential constrictions in the track. Variant E features single stranded scaffold loops at both ends of the docking site. The sequences are complementary to single stranded overhangs at the barrel binding site. The dimerization reaction occurs at 30°C and 3 M NaCl (left). The filament assembly resulted in correctly enclosed pistons inside of filaments (right). Mobility experiments were performed upon adding invader strands that bind to the piston, thereby releasing it from its initial binding site by toehold-mediated strand displacement. Piston variant E showed the highest diffusivity and the longest travel range along the track. All TEM micrographs were high-pass filtered (radius: 25 pixels). All scale bars: 100 nm.

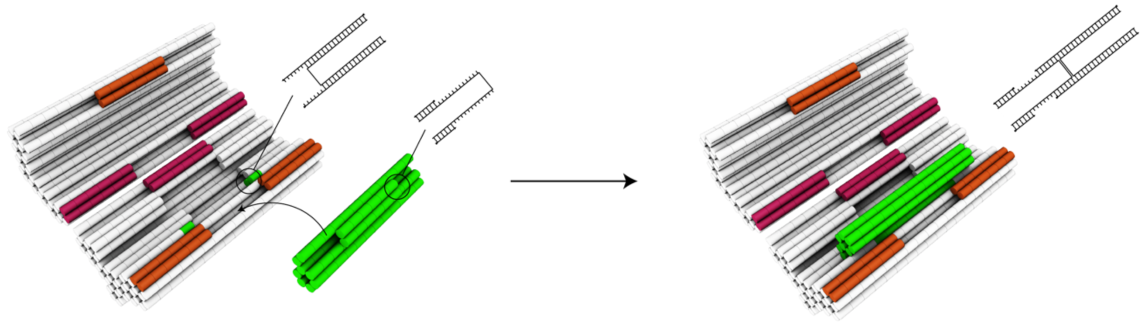

**Figure S 11 Details of the barrel-piston loading reaction.** Models of the barrel (white) and the piston (green) objects. Cylinders indicate DNA double-helices. Orange and magenta cylinders highlight protrusions and recesses involved in closing the barrel. Green cylinders on the inside wall of the barrel object highlights the binding spot for the piston object. Black straight lines pointing away from the black circles highlight zoom-ins on a recess of the barrel (binding spot) and a protrusion on the piston. Zoom-ins: Long black lines indicate the backbone of DNA; shorter black lines indicate bases of DNA. The binding spot on the barrel consists of overall 2 x 6 and 2 x 4 bases long single-stranded overhangs, two on each side (here only two are highlighted on one side). The zoom-in on the piston object highlights the single-stranded scaffold loop at one end of the protrusion (here only one side is highlighted, the other side of the protrusion follows the same design principle).

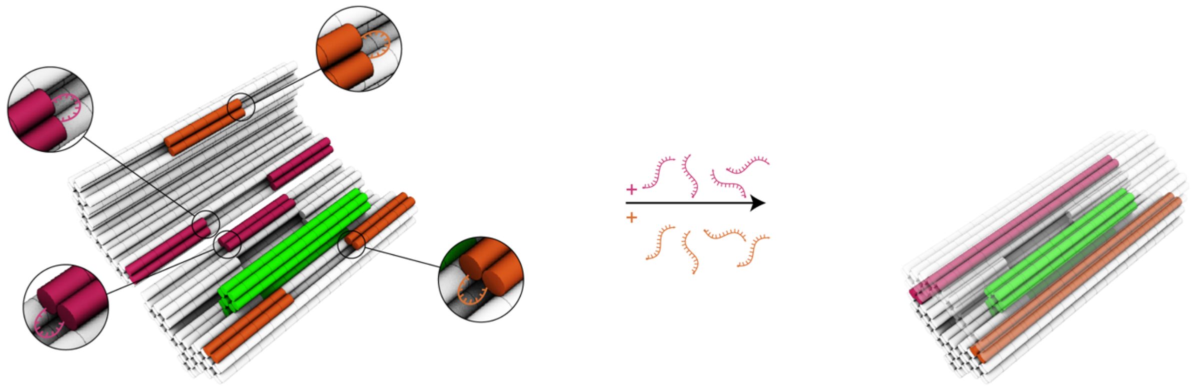

**Figure S 12 Barrel closing mechanism. Left:** Model of a barrel (white)-piston (green) dimer object in an open confirmation. The ends of the orange and magenta protrusions and recesses are initially left single-stranded (insets). **Right:** Oligonucleotides are added in solution and hybridize to the orange and magenta single-stranded loops, thus fixing the barrel in a closed conformation.

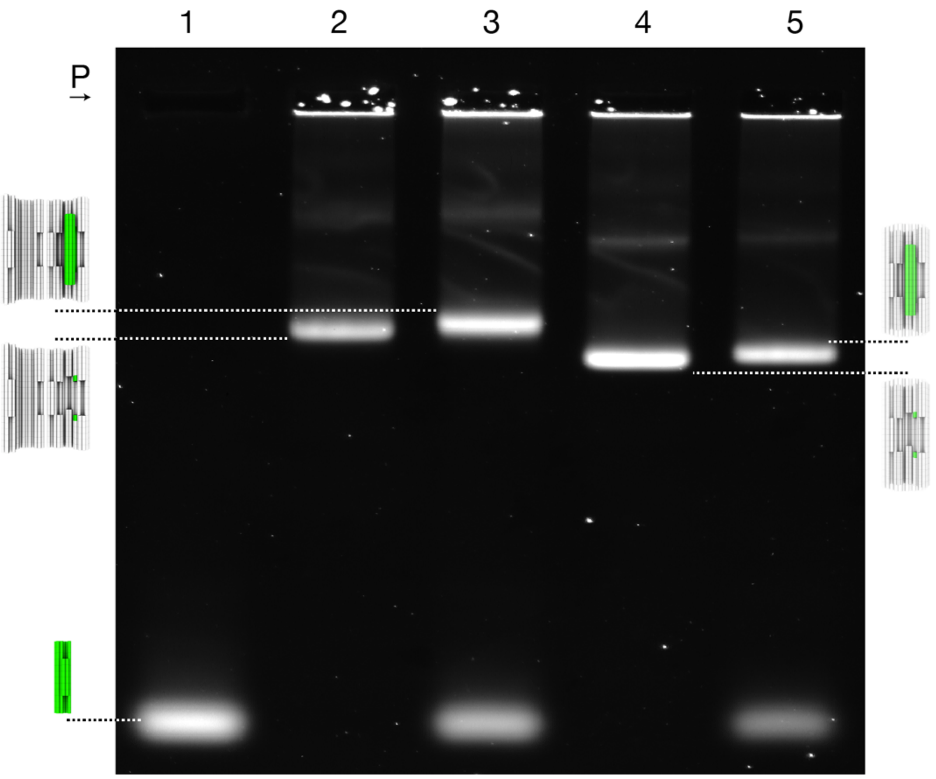


**Figure S 13 EMA quality control of barrel-piston dimers.** Laser-scanned image of a 2.5% agarose gel with 21 mM $MgCl_2$ run on an ice-water bath at 70 V for 180 min on which the following samples were electrophoresed: 1, piston; 2, open barrel; 3, open piston-barrel dimer; 4, closed barrel; 5, closed piston-barrel dimer. Dashed lines point from the corresponding bands to models of the assembled structures.

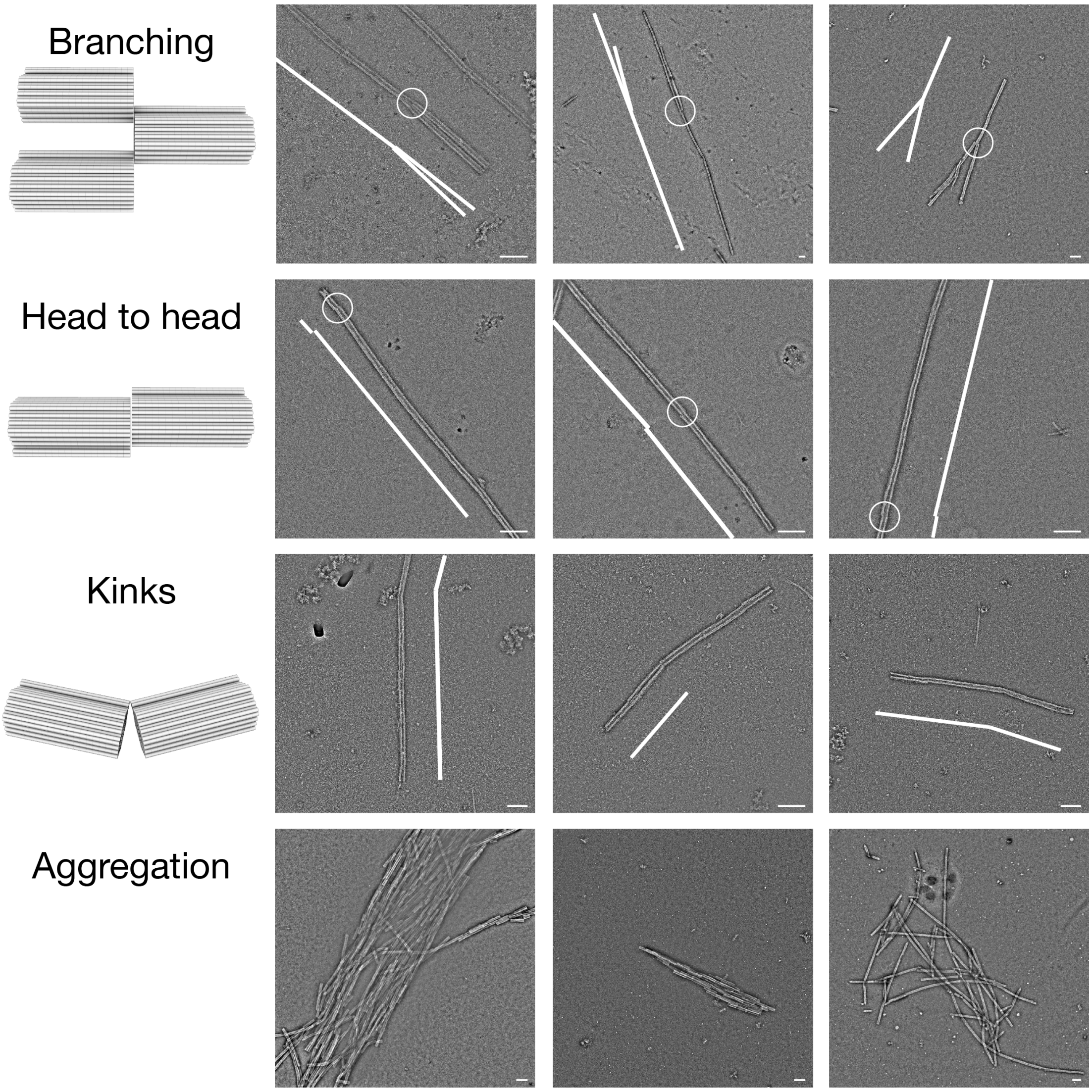


**Figure S 14 Filament defect types. Left:** models of barrels that are attached to each other in wrong orientations. **Right:** exemplary negative-staining TEM images of filaments with defects that represent roadblocks to piston movement. To reduce the occurrence of branching we use two types of polymerization oligonucleotides, one which generate sticky end overhangs and another which produce blunt-ended stacking sites. Sticky end oligonucleotides partly bind to scaffold loops on one end of a barrel and partly on the opposite end of another barrel, thereby bridging two barrels. Stacking end oligonucleotides only bind to scaffold loops on one end of a barrel, resulting in a blunt ended interface. Sticky ends were placed on all helices of the inner layer of the barrel, stacking ends were placed on all helices of the outer layer of the barrel. To reduce head-to-head or tail-to-tail polymerization we implemented a two-step polymerization procedure (Figure S15). Occurrence of kinks could be reduced by using a polymerization oligonucleotide to barrel stoichiometry of no greater than 4:1. The tendency to form aggregates was least when polymerization was performed in the presence of 2 M NaCl at 40°C out of all conditions tested. Using an additional incubation step of 1 hour at 40°C and 2 M NaCl after closing assembled barrel-piston dimers and right before triggering the polymerization reaction by adding the polymerization oligonucleotides was also beneficial. Images were acquired with a Philips

CM100 microscope. All TEM micrographs were high-pass filtered (radius: 25 pixels). All scale bars: 100 nm.

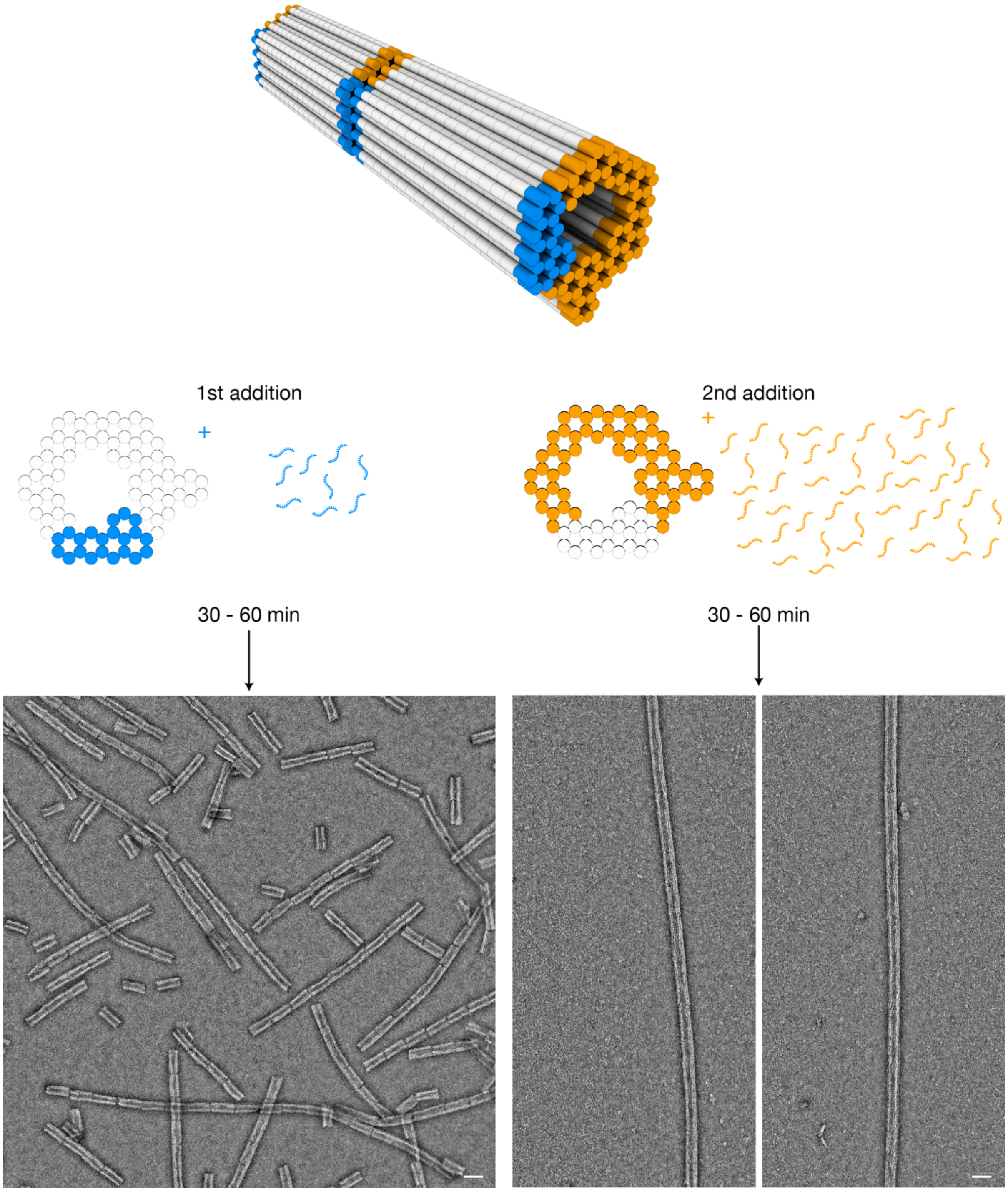


**Figure S 15 Best polymerization strategy**. **Top:** models of 2 barrels. White, blue and orange cylinders represent DNA double-helices. **Middle:** cross sectional view of the barrel. **Left:** first addition of oligonucleotides that serve as a first set of polymerization oligonucleotides. They bind to the scaffold loops at the end of 1 (blue circles) of 6 sides of the barrel. **Bottom left:** typical field of view negative-staining TEM micrograph of barrels 30-60 min after adding the first set of polymerization oligonucleotides and incubating at 40°C and 2 M NaCl, imaged with a Philips CM100 microscope. **Middle right:** second addition of oligonucleotides that serve as a second set of polymerization oligonucleotides. They bind to the scaffold loops at the end of 5 (orange circles) of 6 sides of the barrel.

**Bottom right:** typical field of view negative-staining TEM micrograph of barrels 30-60 min after adding the first set of polymerization oligonucleotides and incubating at 40°C and 2 M NaCl and then adding the second set of polymerization oligonucleotides and incubating at 40°C and 2 M NaCl, imaged with a Philips CM100 microscope. All TEM micrographs were high-pass filtered (radius: 25 pixels). Scale bars: 50 nm.

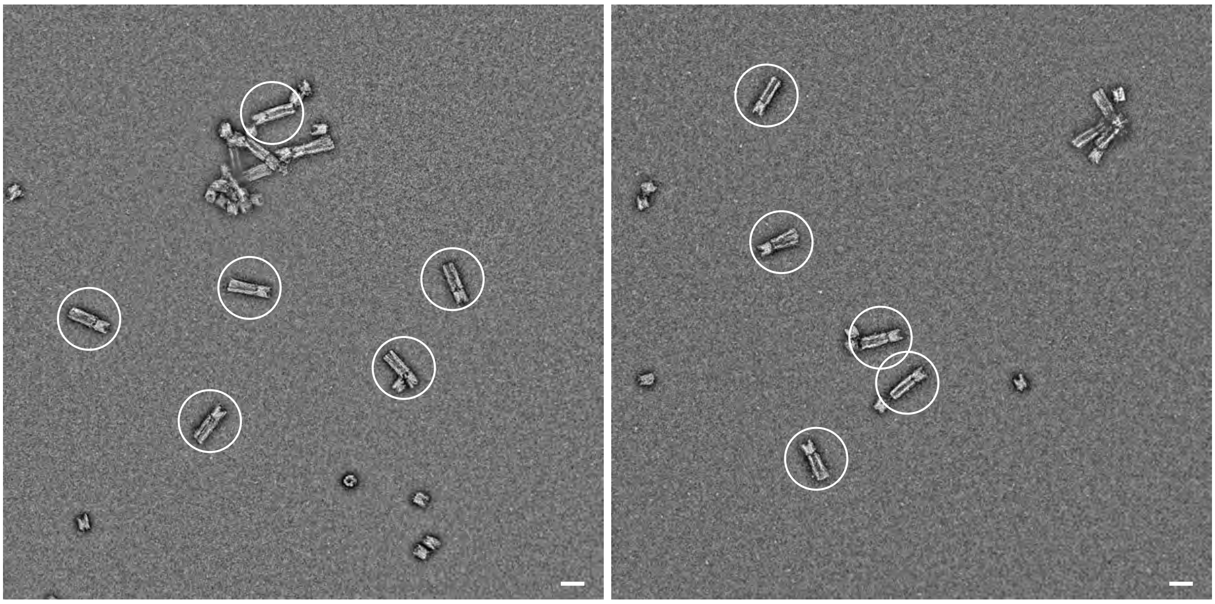

**Figure S 16 TEM quality control of cap1-barrel dimers.** Typical field of view negative-staining TEM micrographs of assembled barrel-cap1 dimers, imaged with a Philips CM100 microscope. The TEM micrographs were high-pass filtered (radius: 25 pixels). Scale bars: 50 nm.

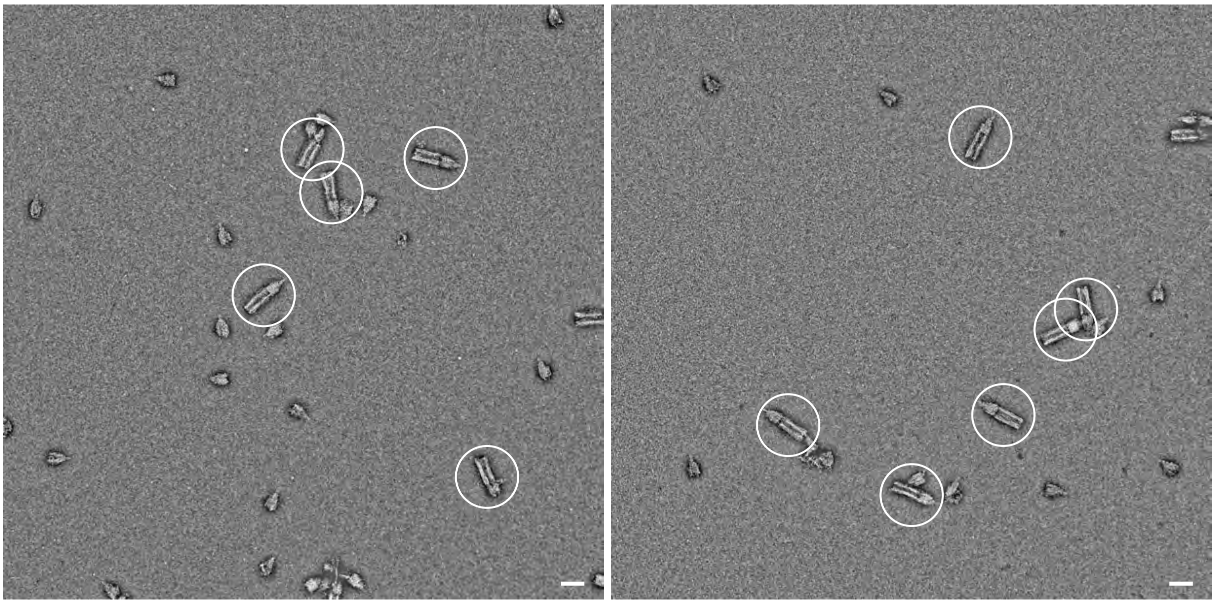

**Figure S 17 TEM quality control of cap2-barrel dimers.** Typical field of view negative-staining TEM micrographs of assembled barrel-cap2 dimers, imaged with a Philips CM100 microscope. The TEM micrographs were high-pass filtered (radius: 25 pixels). Scale bars: 50 nm.

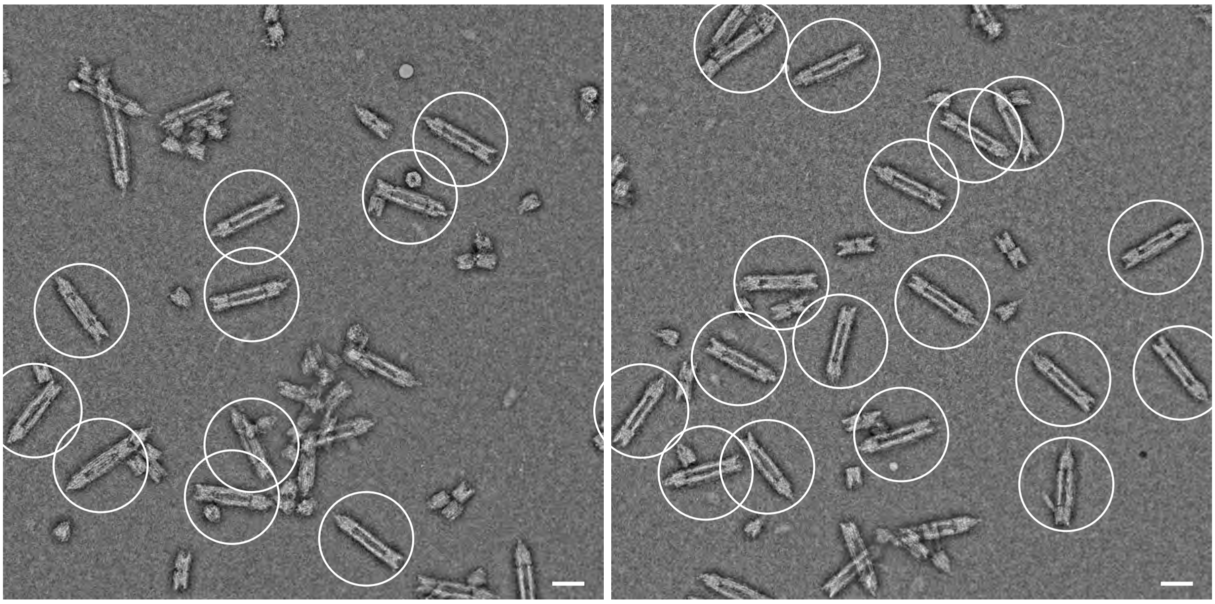

**Figure S 18 TEM quality control of cap1-barrel-cap2 trimers.** Typical field of view negative-staining TEM micrographs of assembled cap1-barrel-cap2 trimers, imaged with a Philips CM100 microscope. The TEM micrographs were high-pass filtered (radius: 25 pixels). Scale bars: 50 nm.

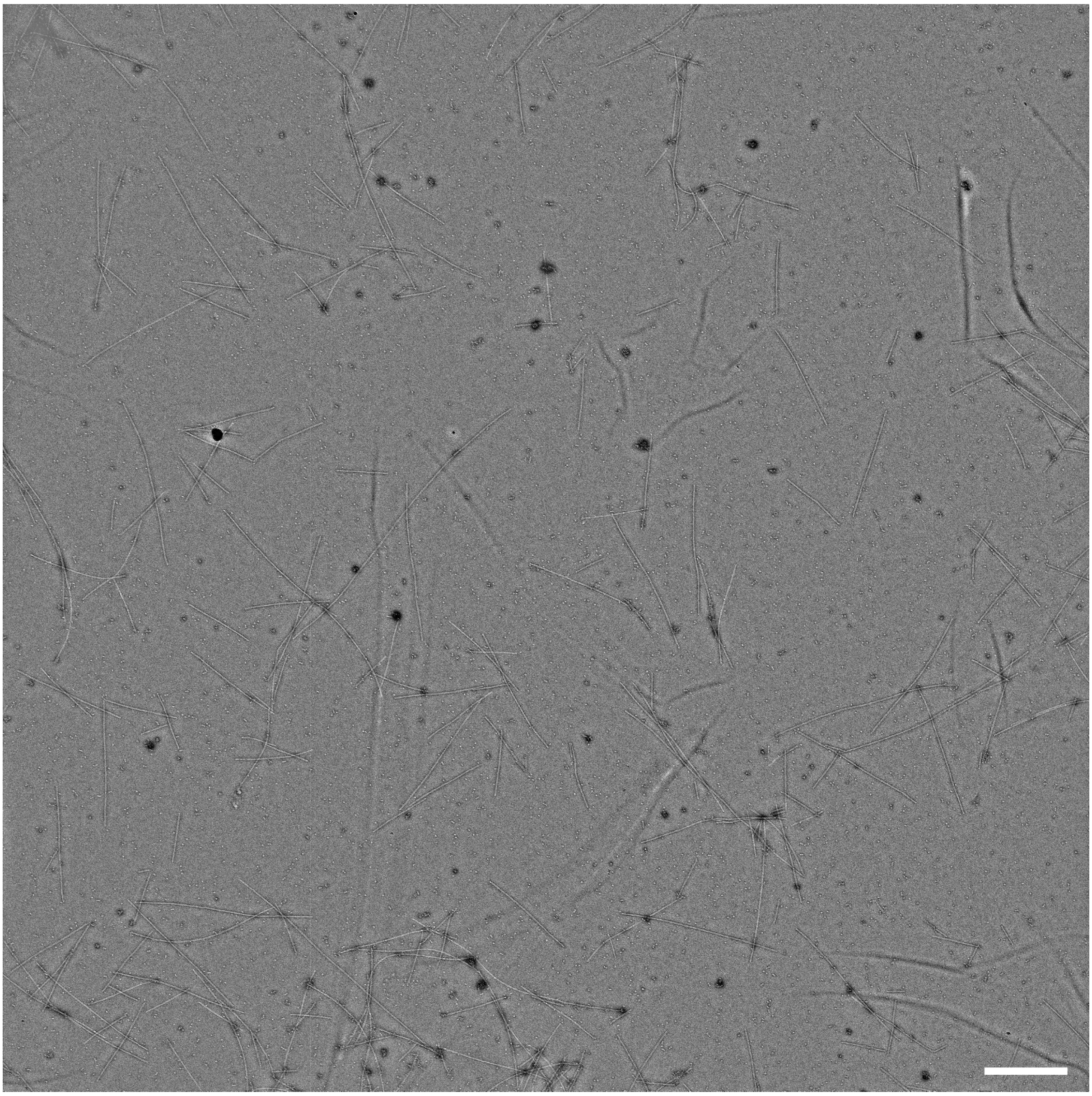

**Figure S 19 TEM quality control of assembled tracks containing pistons.** Typical field of view negative-staining TEM micrograph of assembled transport systems imaged with a Jeol JEM3200 FSC microscope. The TEM micrograph was high-pass filtered (radius: 50 pixels). Scale bar: 1 μm.

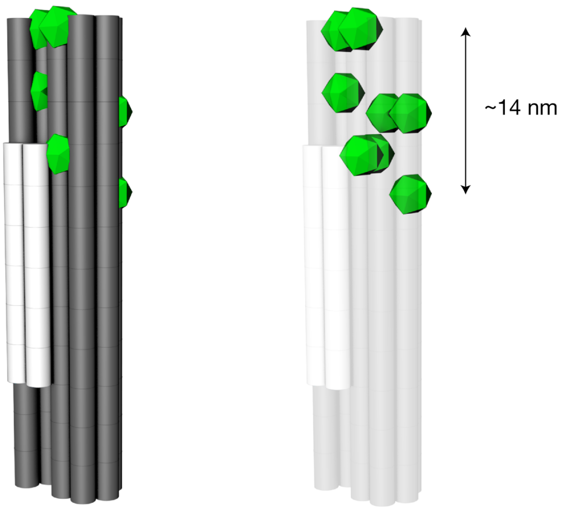


**Figure S 20 Fluorescent labeling of the piston.** Illustration of the Cyanine-3 dye placement on the piston. Cylinders represent DNA double-helices. Green objects illustrate Cyanine-3 dyes. The 8 Cyanine-3 dyes are placed over a distance of 14 nm.

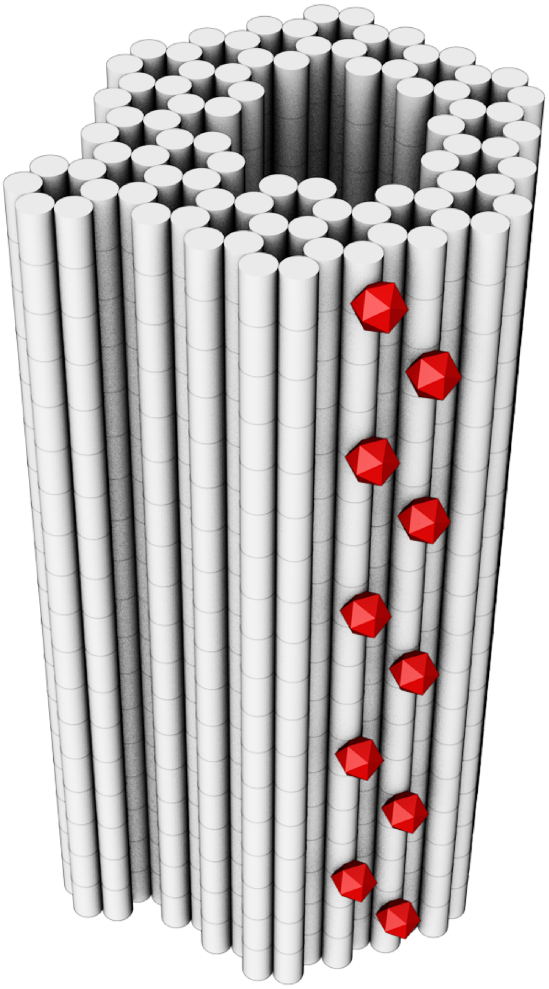

**Figure S 21 Fluorescent labelling of the barrel.** Illustration of the Cyanine-5 dye placement on the barrel. Cylinders represent DNA double-helices. Red objects illustrate Cyanine-5 dyes.

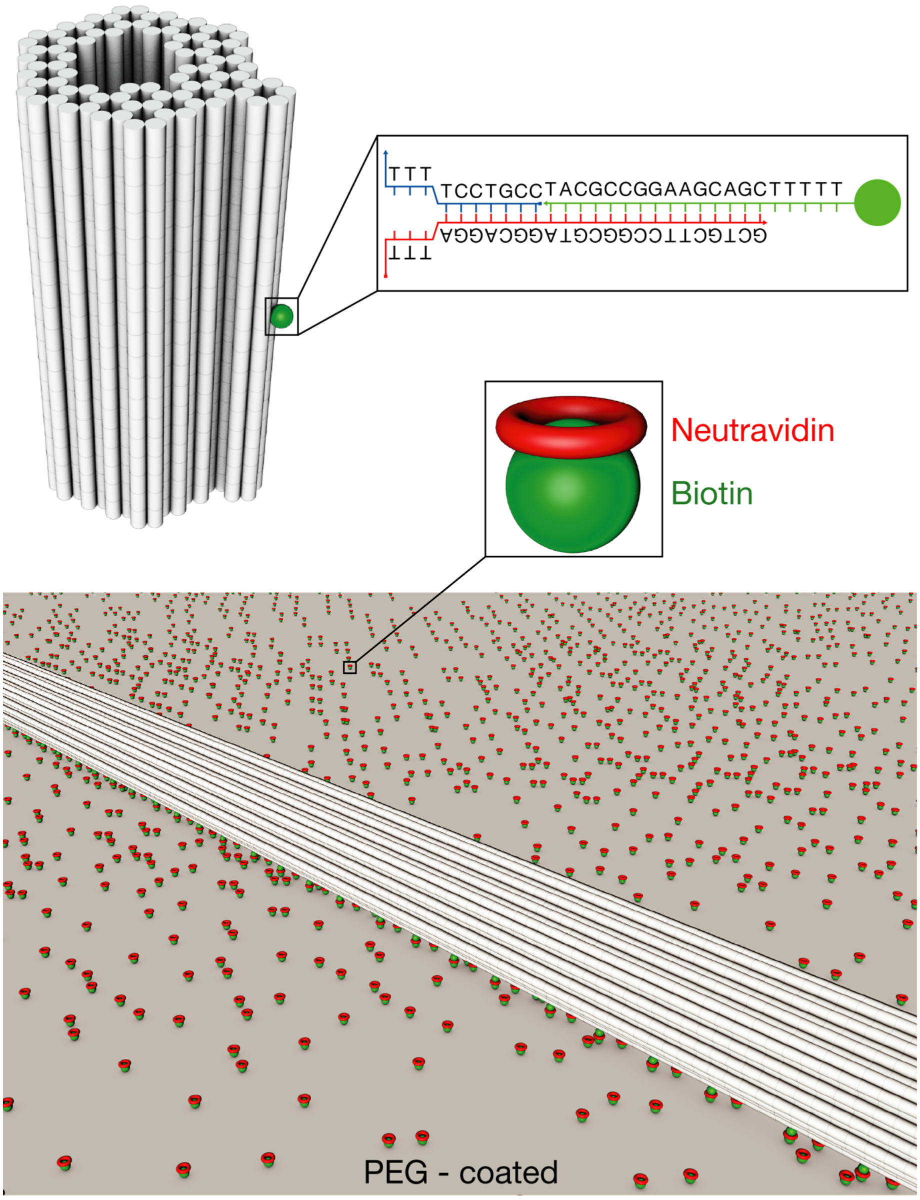


**Figure S 22 Biotin placement on the barrel.** Illustration of the biotin placement on the barrel. **Top left:** Cylinders represent DNA double-helices. Green sphere represents biotin. **Top right:** Zoom-in on the biotin anchor. Letters indicate the sequences, straight lines represent single bases and backbones of three (red, blue, green) oligonucleotides. **Bottom:** Illustration of a tunnel bound to a pegylated glass surface via neutravidin. Grey surface represents the pegylated glass surface (protocol in materials & methods). White cylinders represent DNA double-helices. Green spheres represent biotin, red objects represent neutravidin. Zoom-in highlights a biotin on the glass surface, bound to one neutravidin.

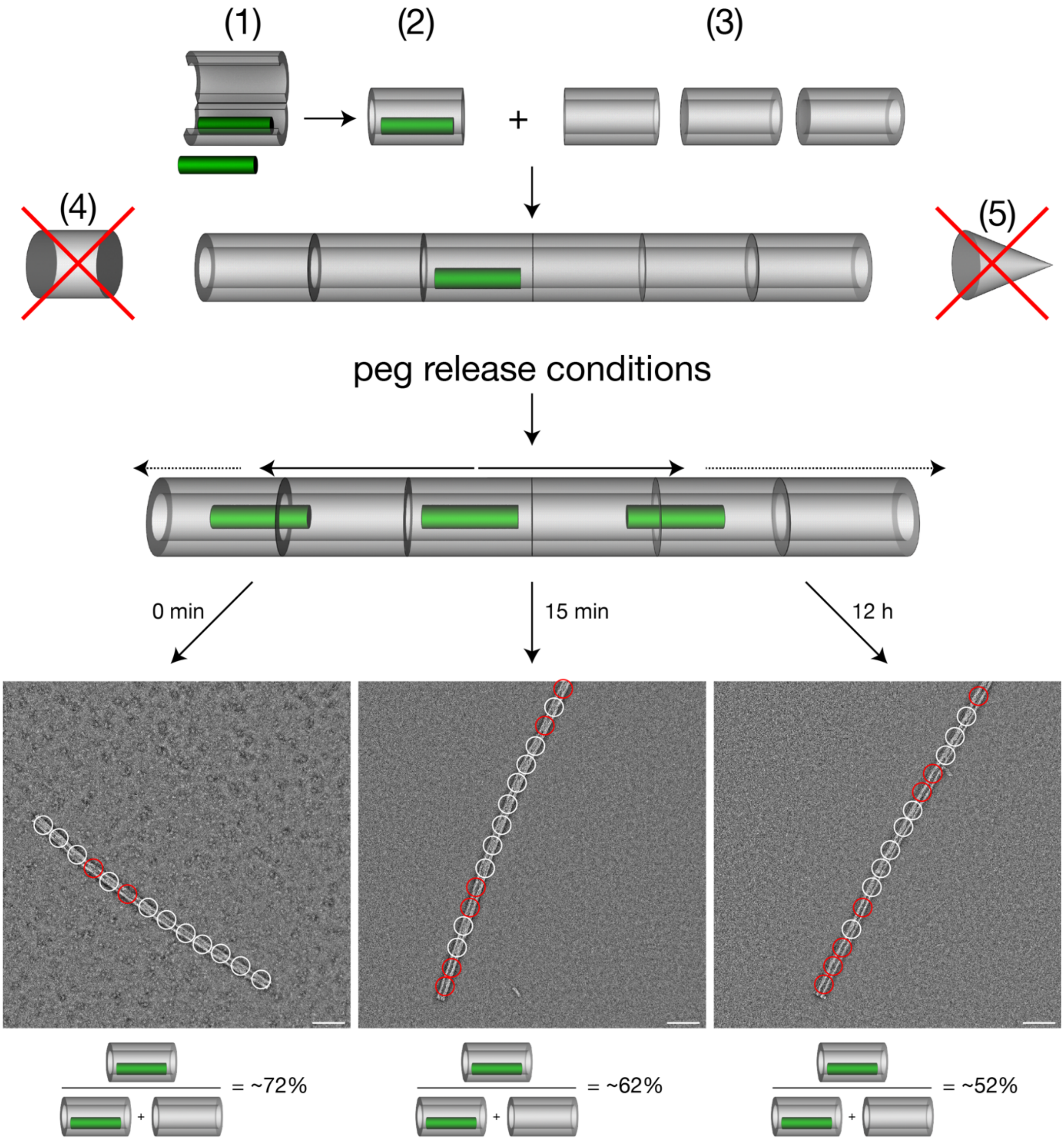


**Figure S 23 TEM "efflux" analysis of low mobility piston-track designs. Top:** Schematic illustration of an experiment designed to test very low mobility pistons for their functionality. Grey cylinders represent the barrel, green objects represent the piston. For this experiment, filaments with pistons inside but without any caps attached at the ends are needed. To this end, the piston variant in question was bound to the barrel at barrel to piston ratios between 1:1 and 2:1 **(1)**. The barrel was then subsequently closed permanently **(2)**. Polymerization oligonucleotides were then added to this sample, containing barrel-piston dimers and empty barrels **(3)**. Steps **(4)** and **(5)**, adding both caps to close the filament ends, were omitted, resulting in open filaments, with different amounts of pistons inside, depending on the initial barrel to piston ratio. These open filament samples were then split into separate reaction tubes and subjected to different piston releasing conditions, based on the specific piston variant in question, and incubated at these conditions for various lengths of time. The resulting samples were then analyzed using negative-staining TEM (bottom), by counting the number of vacant (highlighted by red circles) and occupied (highlighted by white circles) piston binding spots inside of filaments. **Bottom:** 3 exemplary negative-staining TEM micrographs of such samples at different points in time and the respective fraction of occupied binding spots and total number of binding spots (below TEM micrographs). The TEM micrographs were high-pass filtered (radius: 25 pixels). Scale bars: 100 nm. Several hundred binding spots were analyzed and counted for all samples. **Bottom left:** this specific sample was not subjected to piston releasing conditions and showed ~72% occupied binding spots. **Bottom middle:** this sample, initially from the same pool as the first sample (left), incubated at piston releasing conditions for 15 min before TEM grids were prepared. It shows ~62% occupied binding spots. **Bottom right:** this sample incubated at piston releasing conditions for 24 hours before TEM grids were prepared. It shows ~52% occupied binding spots.

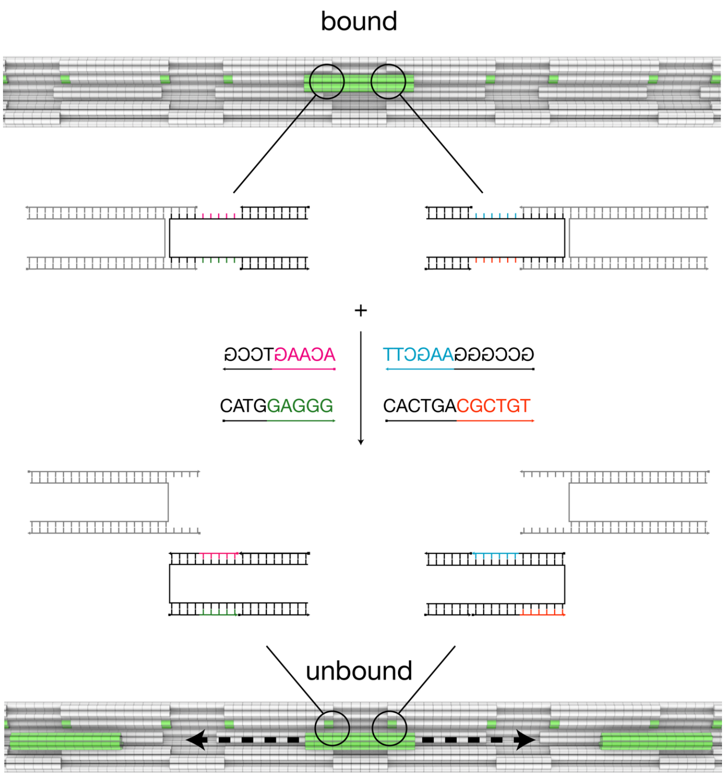


**Figure S 24 Illustration of the piston release mechanism.** Cylinders represent DNA double-helices. **Top:** snippet of a filament with a piston (green object) trapped inside of the filament, bound in its initial binding spot. Zoom-ins (black circles) highlight where and how the piston is initially bound. Black, magenta, blue, green and orange lines indicate the backbone of single and double stranded DNA. Short lines indicate individual bases of the DNA. Letters indicate the sequences of invader strands. Upon addition of the invader strands **(middle)** the piston is released from its initial binding spot by toehold-mediated strand displacement. The invader strands bind to their respective positions on the piston, defined by their sequences. The piston is then free to move back and forth along the direction of the filament **(bottom)**, indicated by the dashed black arrows.

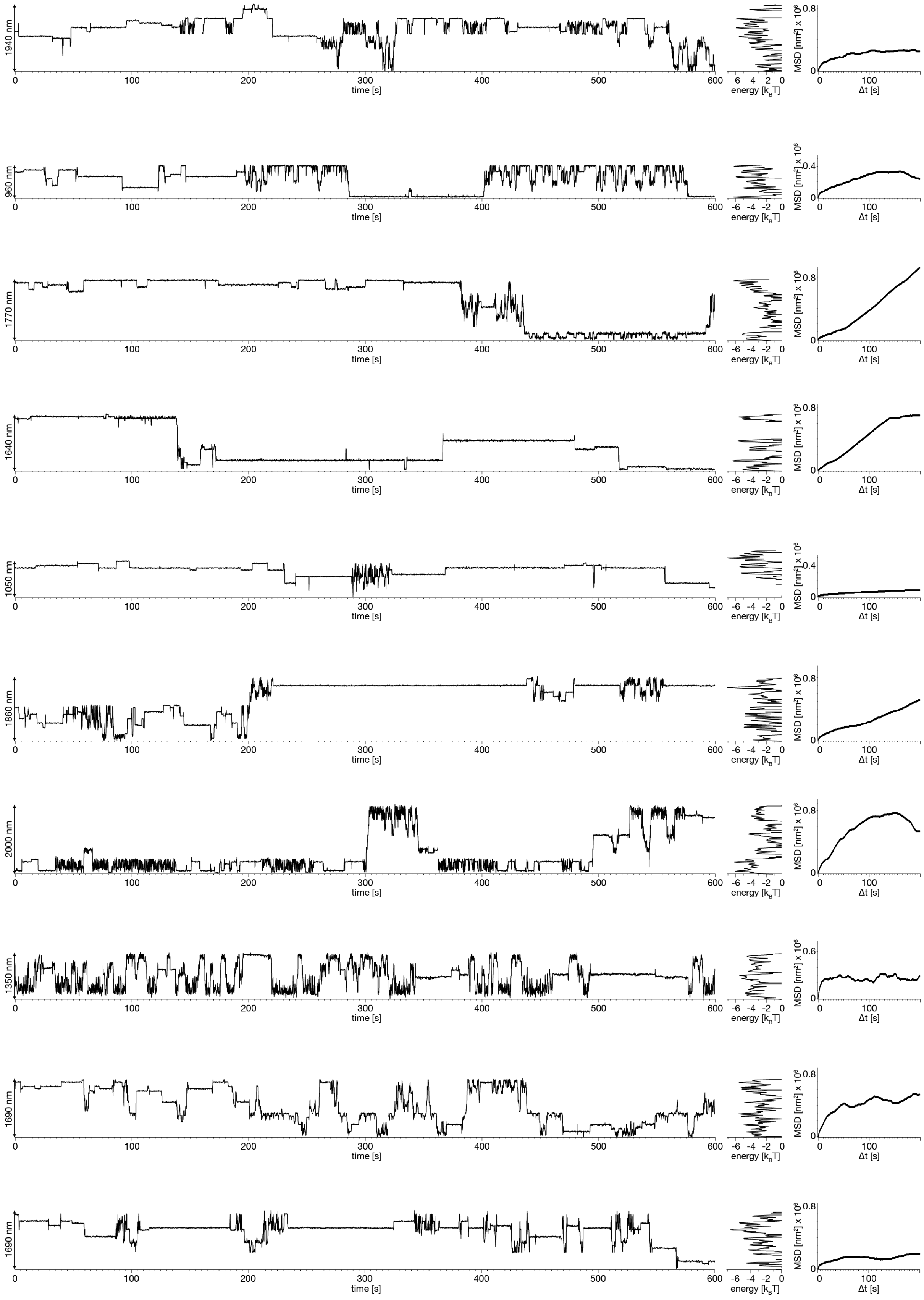


**Figure S 25** Left: Position-time traces of individual pistons (1-10) recorded at 20°C ambient temperature. Middle: Energy profiles computed from position probability distributions for the traces. Right: Mean square displacement (MSD) curves of the single particle traces.

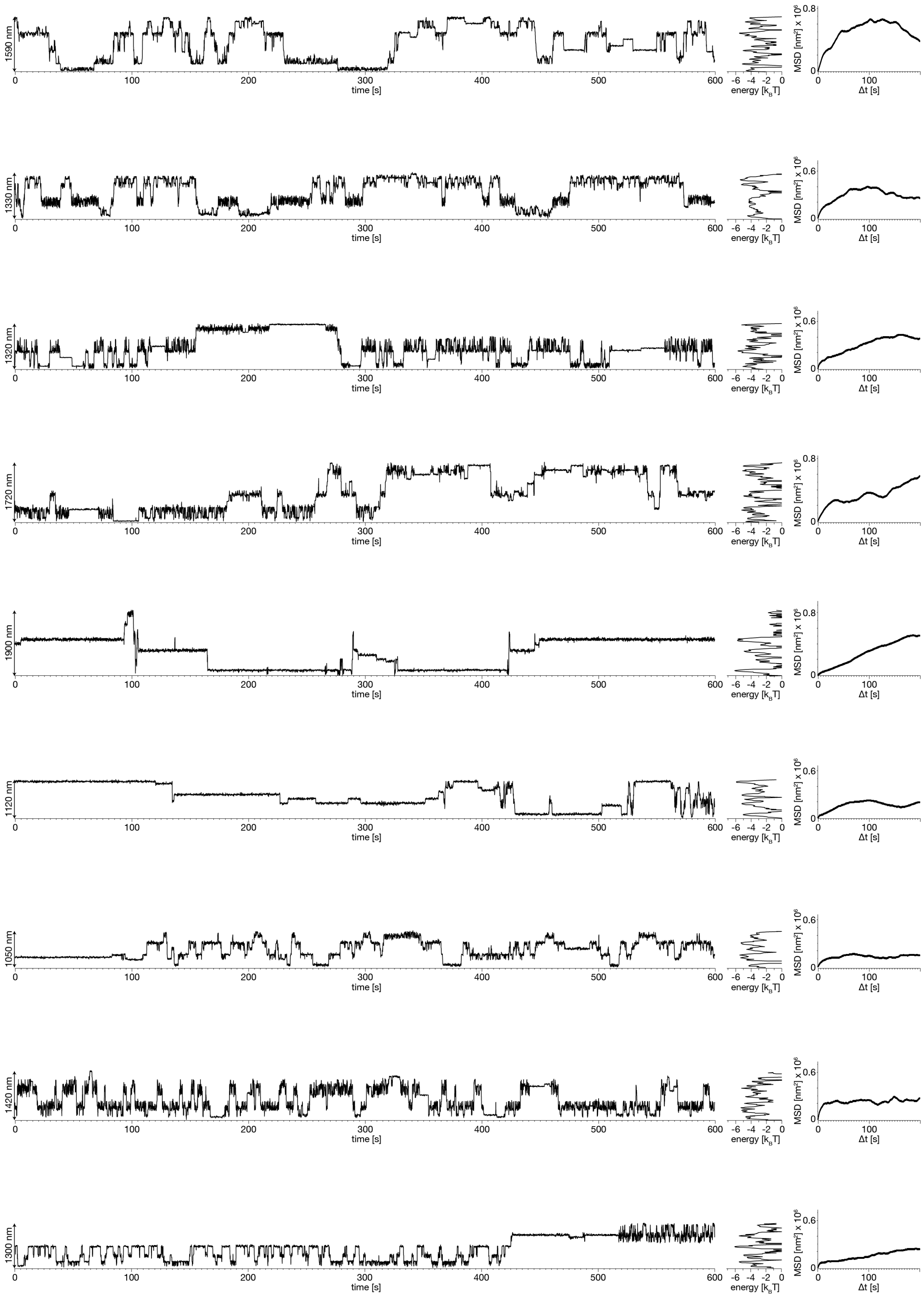


**Figure S 26** Left: Position-time traces of individual pistons (11-19) recorded at 20°C ambient temperature. Middle: Energy profiles computed from position probability distributions for the traces. Right: Mean square displacement (MSD) curves of the single particle traces.

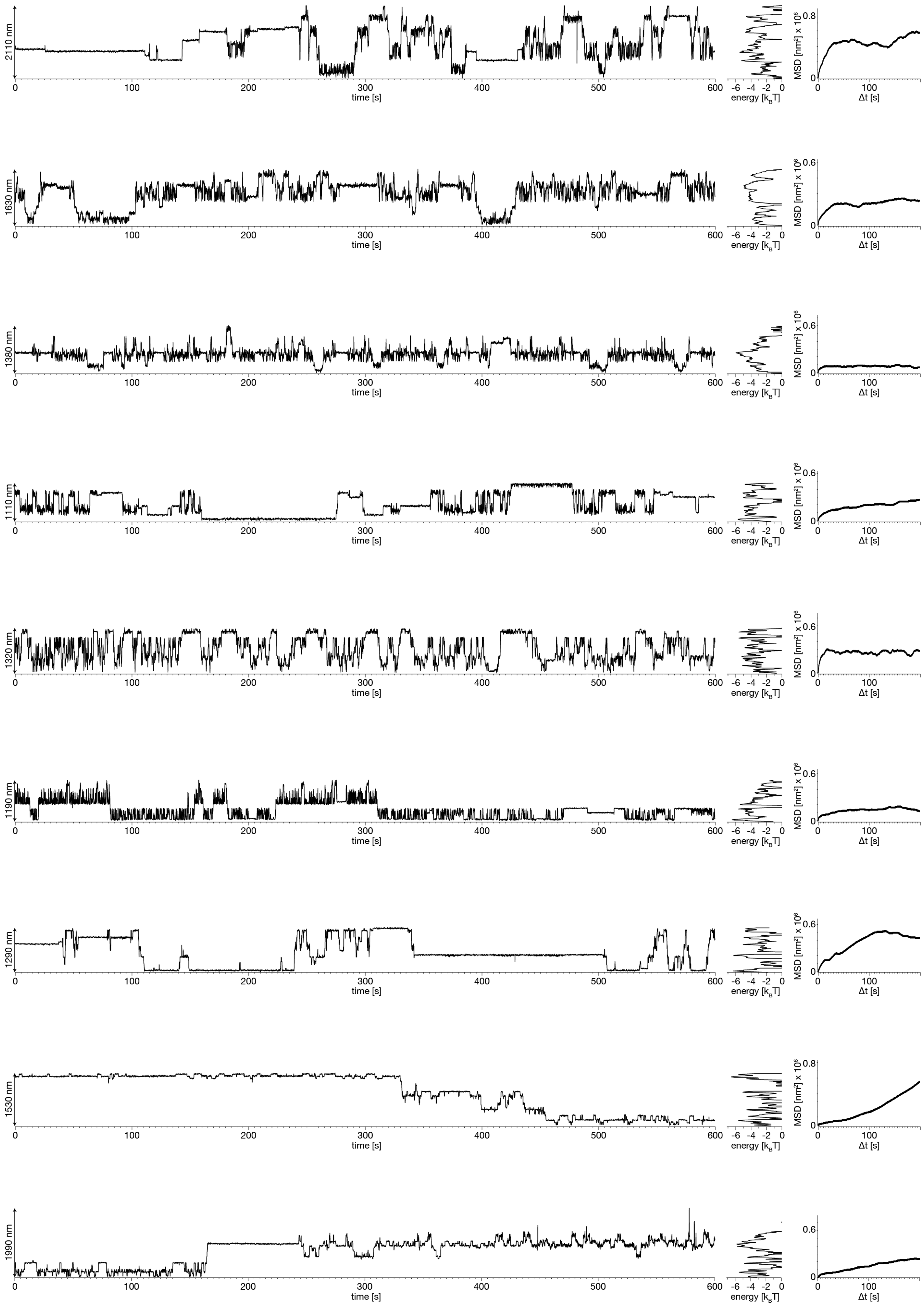


**Figure S 27** Left: Position-time traces of individual pistons (20-28) recorded at 20°C ambient temperature. Middle: Energy profiles computed from position probability distributions for the traces. Right: Mean square displacement (MSD) curves of the single particle traces.

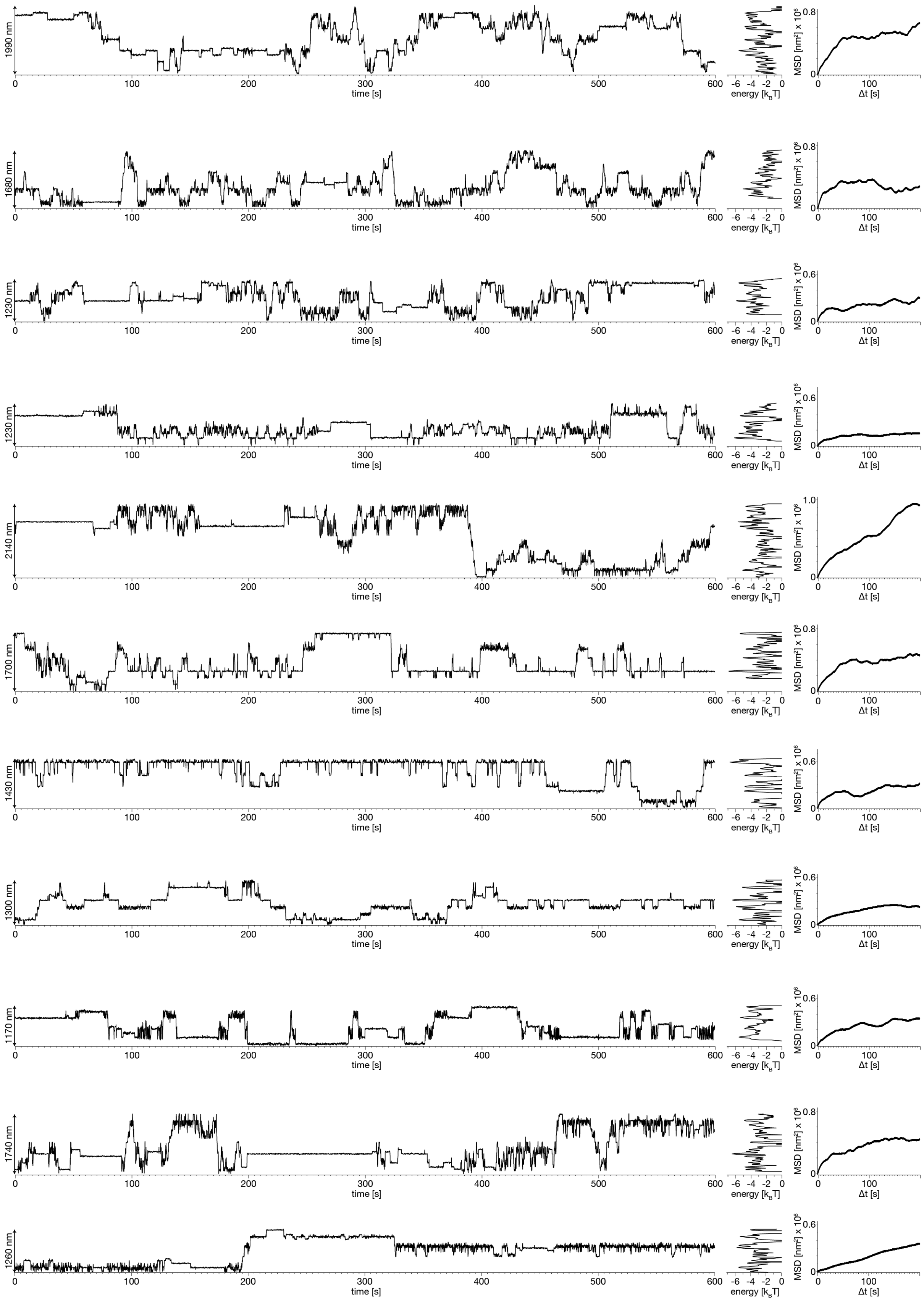


**Figure S 28** Left: Position-time traces of individual pistons (29-39) recorded at 25°C ambient temperature. Middle: Energy profiles computed from position probability distributions for the traces. Right: Mean square displacement (MSD) curves of the single particle traces.

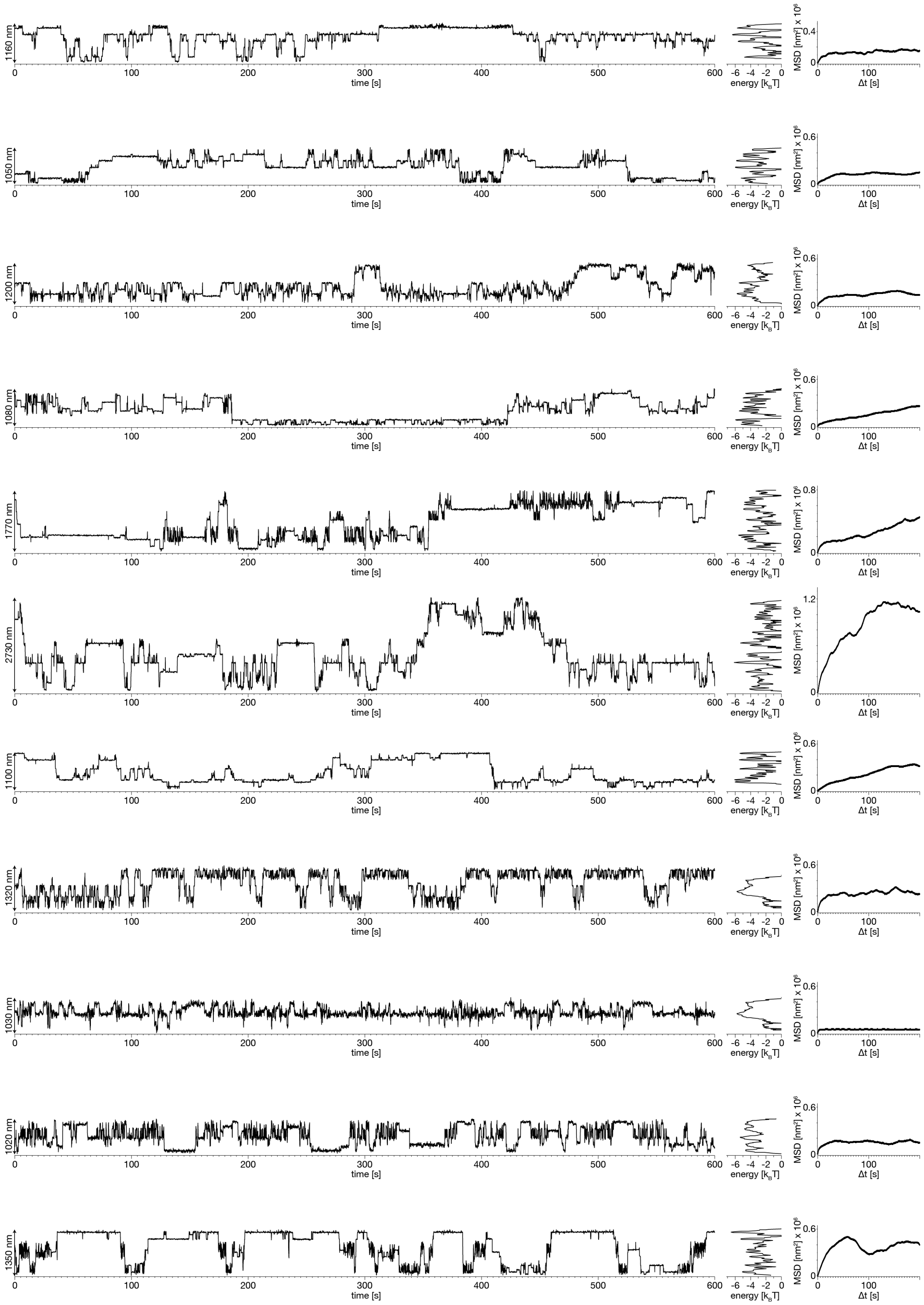


**Figure S 29** Left: Position-time traces of individual pistons (40-50) at 25°C ambient temperature. Middle: Energy profiles computed from position probability distributions for the traces. Right: Mean square displacement (MSD) curves of the single particle traces.

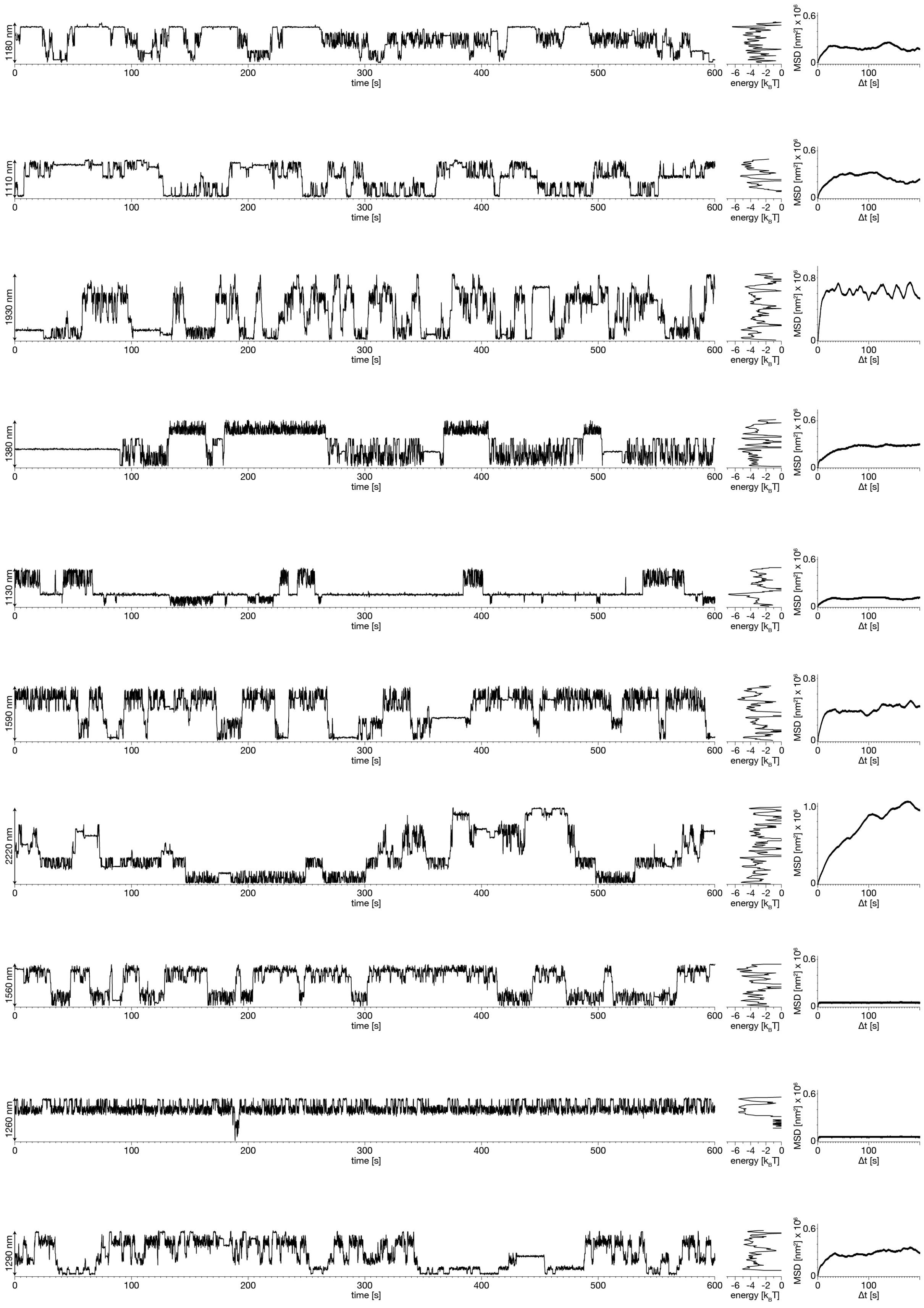


**Figure S 30** Left: Position-time traces of individual pistons (51-60) at 25°C ambient temperature. Middle: Energy profiles computed from position probability distributions for the traces. Right: Mean square displacement (MSD) curves of the single particle traces.

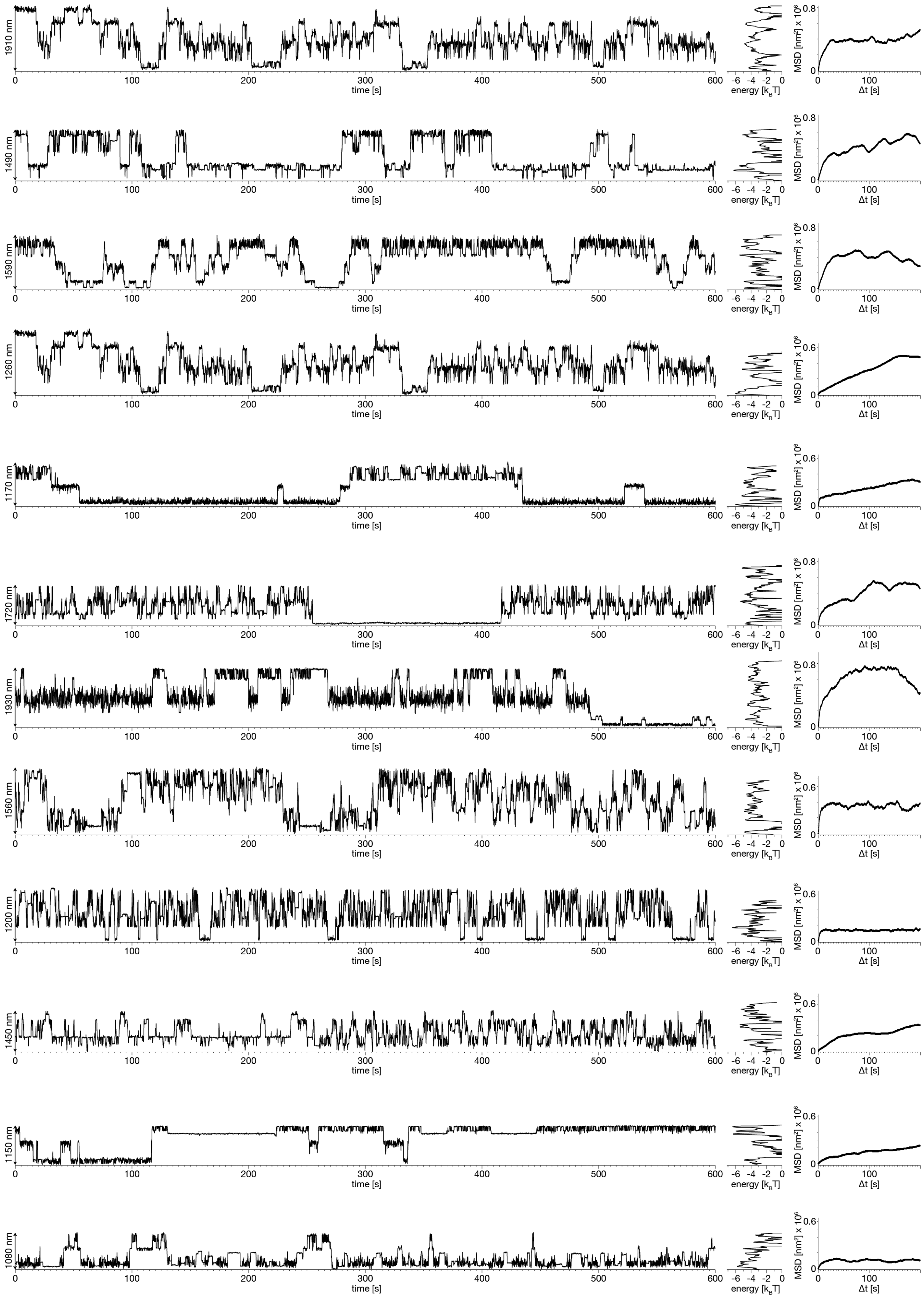


**Figure S 31** Left: Position-time traces of individual pistons (61-72) at 30°C ambient temperature. Middle: Energy profiles computed from position probability distributions for the traces. Right: Mean square displacement (MSD) curves of the single particle traces.

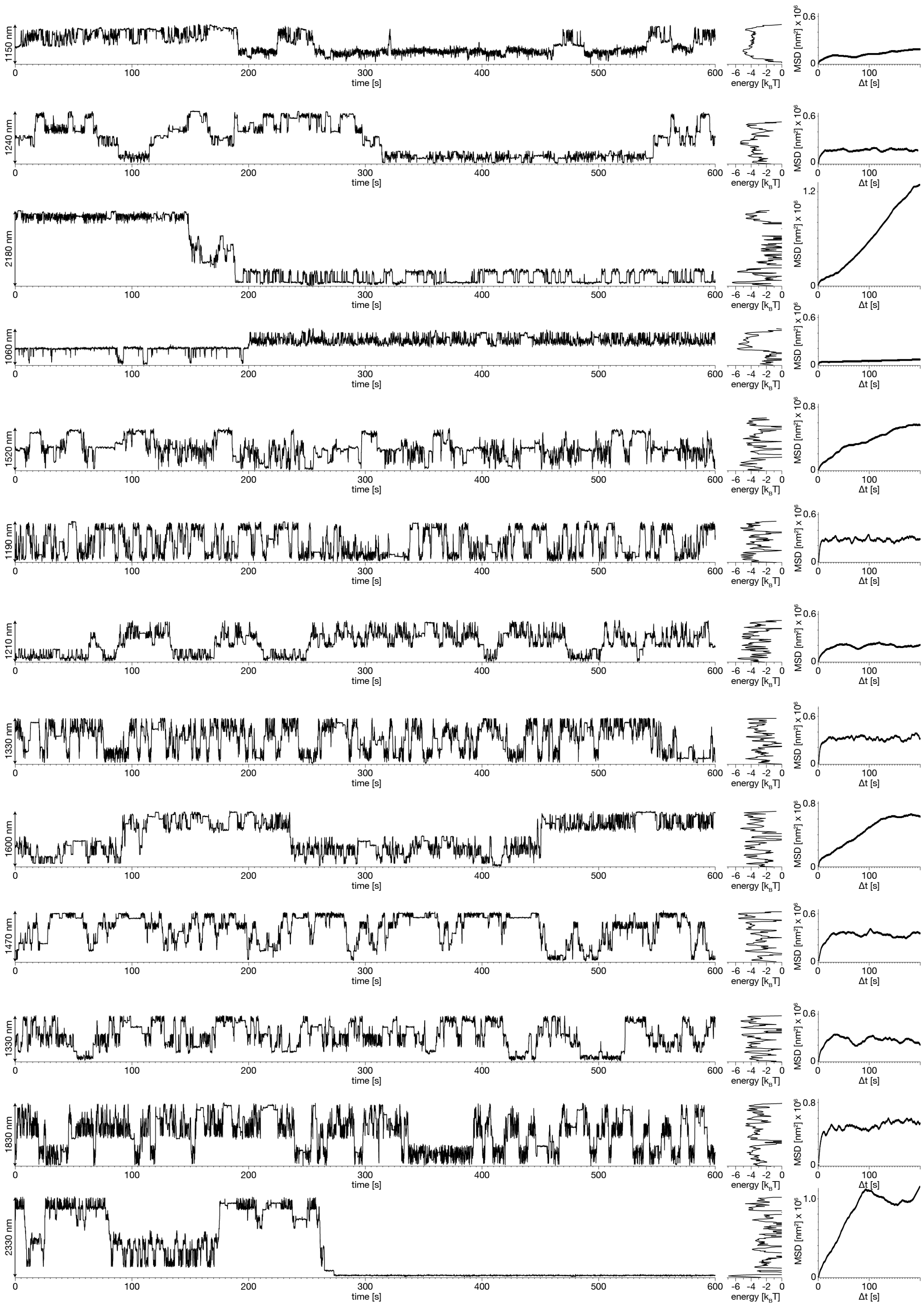


**Figure S 32** Left: Position-time traces of individual pistons (73-85) at 30°C ambient temperature. Middle: Energy profiles computed from position probability distributions for the traces. Right: Mean square displacement (MSD) curves of the single particle traces.

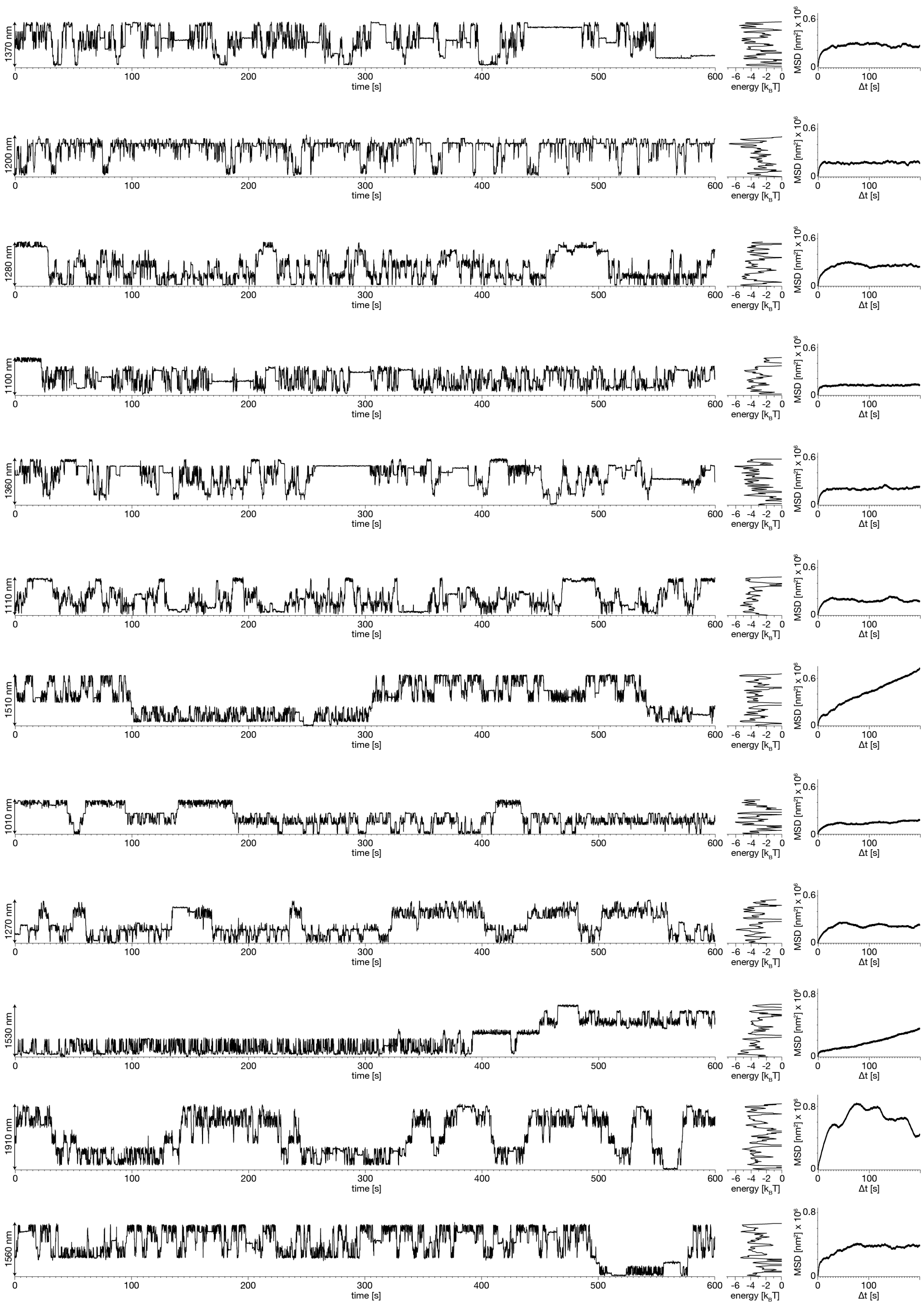


**Figure S 33** Left: Position-time traces of individual pistons (86-97) at 30°C ambient temperature. Middle: Energy profiles computed from position probability distributions for the traces. Right: Mean square displacement (MSD) curves of the single particle traces.

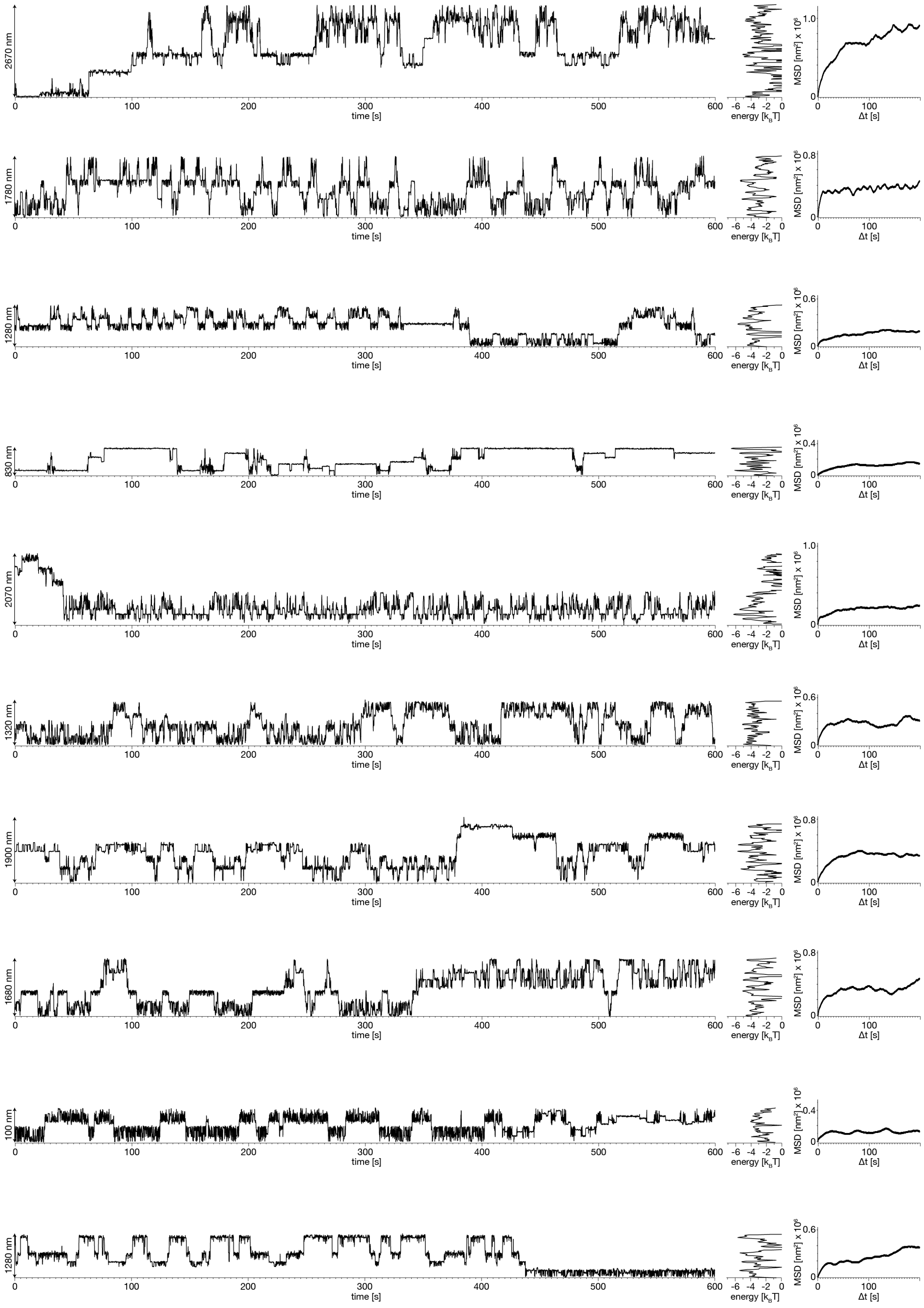


**Figure S 34** Left: Position-time traces of individual pistons (98-107) at 35°C ambient temperature. Middle: Energy profiles computed from position probability distributions for the traces. Right: Mean square displacement (MSD) curves of the single particle traces.

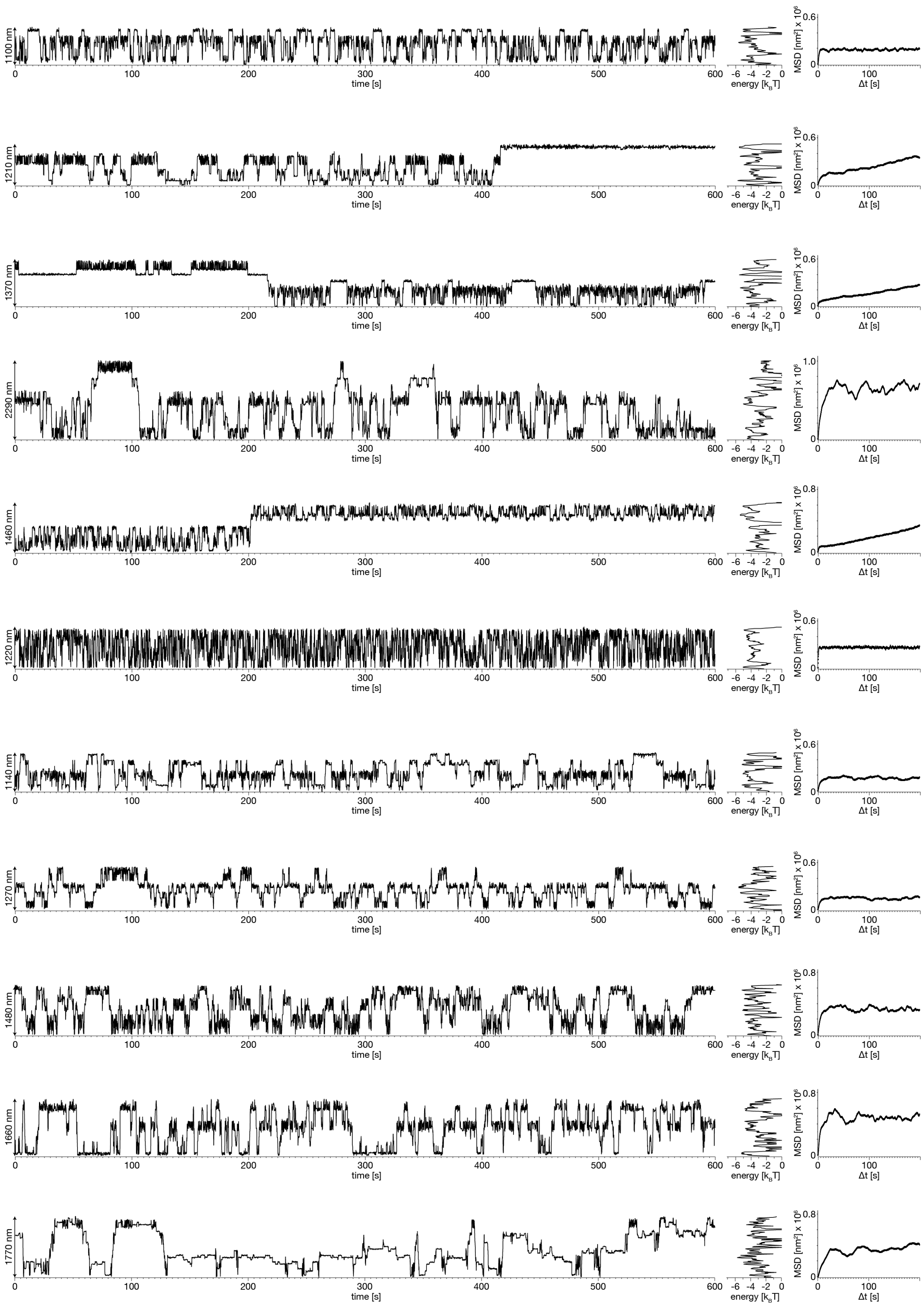


**Figure S 35** Left: Position-time traces of individual pistons (108-118) at 35°C ambient temperature. Middle: Energy profiles computed from position probability distributions for the traces. Right: Mean square displacement (MSD) curves of the single particle traces.

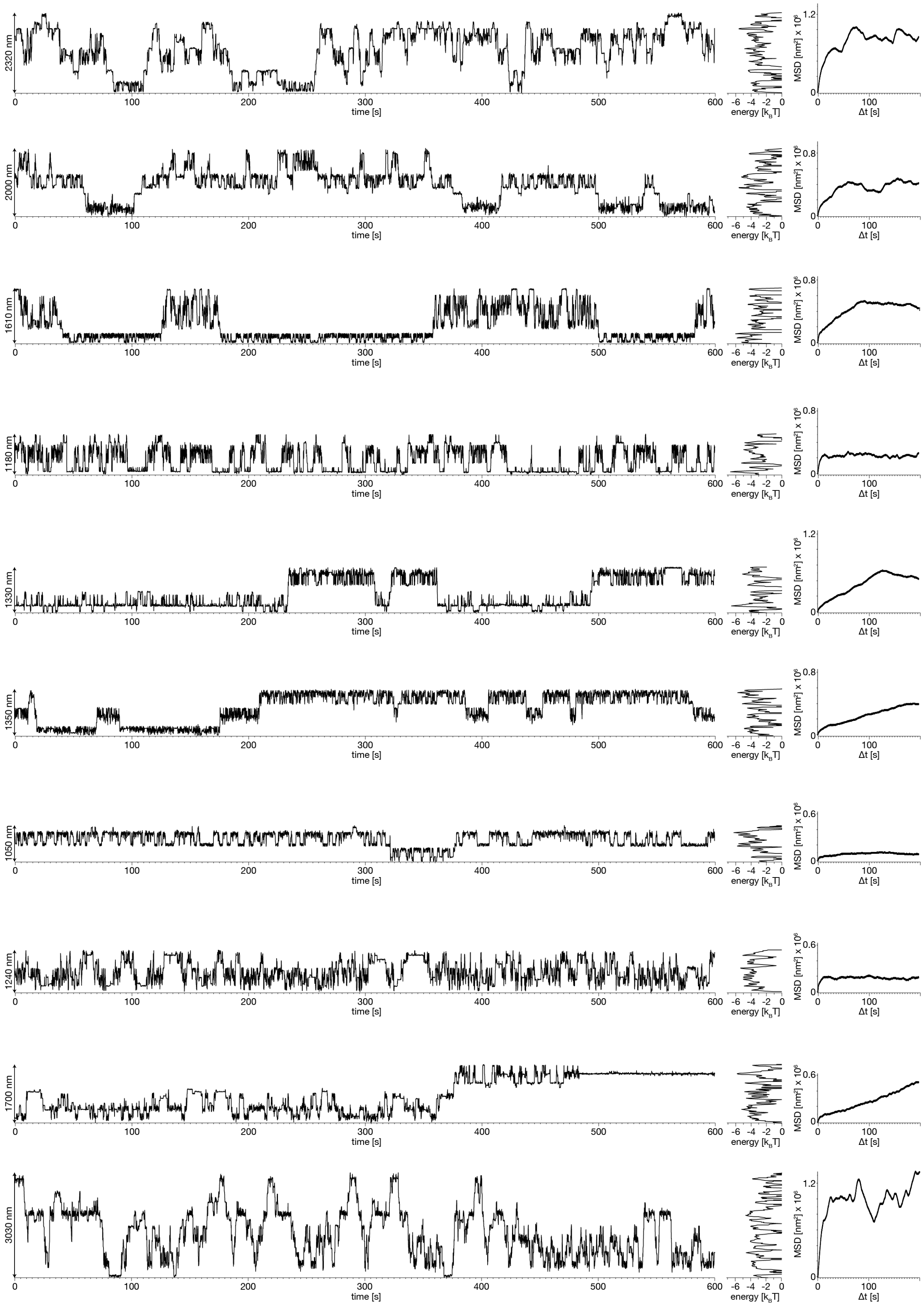


**Figure S 36** Left: Position-time traces of individual pistons (119-128) at 35°C ambient temperature. Middle: Energy profiles computed from position probability distributions for the traces. Right: Mean square displacement (MSD) curves of the single particle traces.

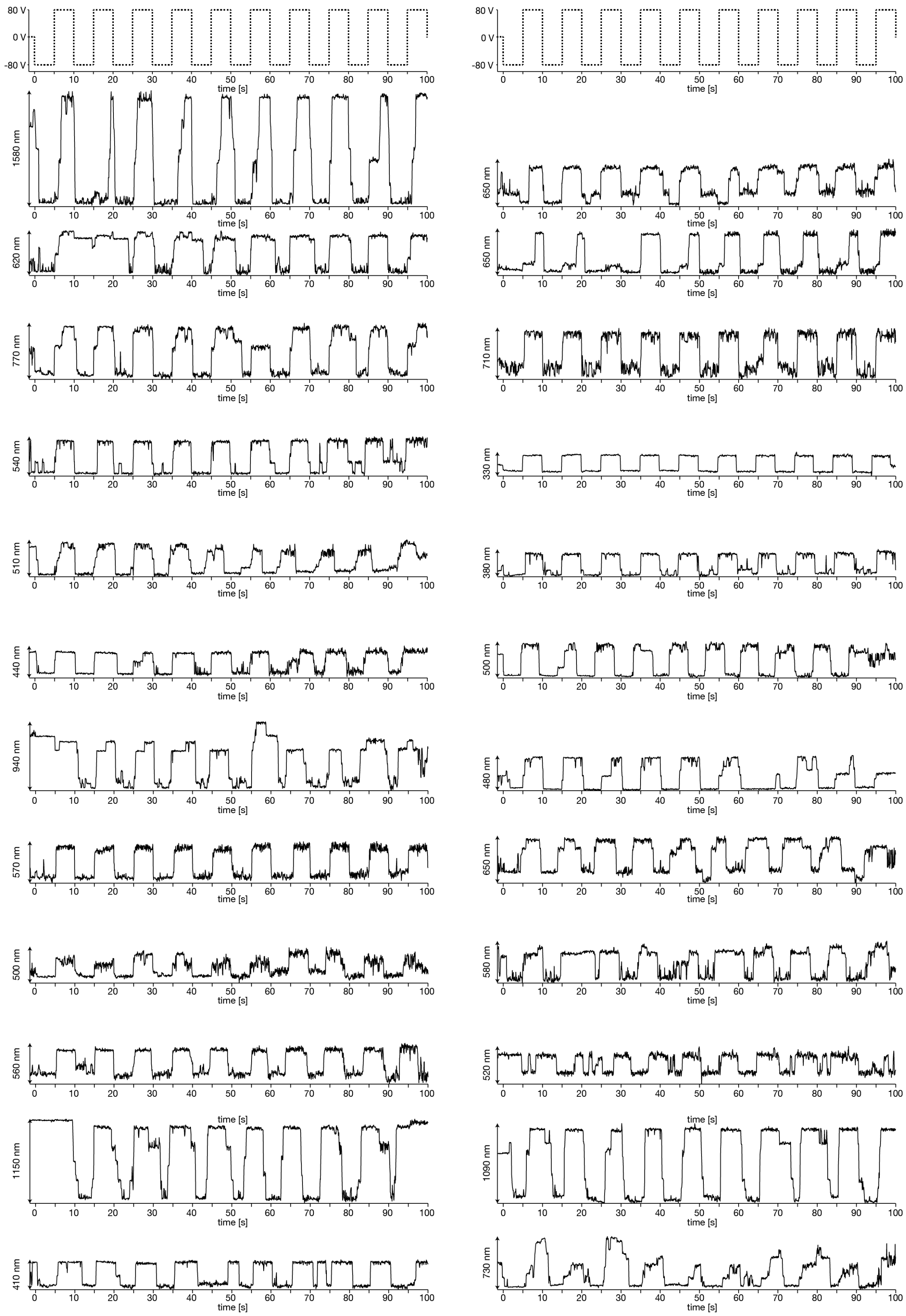


**Figure S 37 Driving the pistons with electric fields.** Position-time traces of 24 pistons moving inside tunnels, with the tunnels oriented at angles between 0-50° relative to an applied electric field that was switched back and forth every 5 seconds (top row).

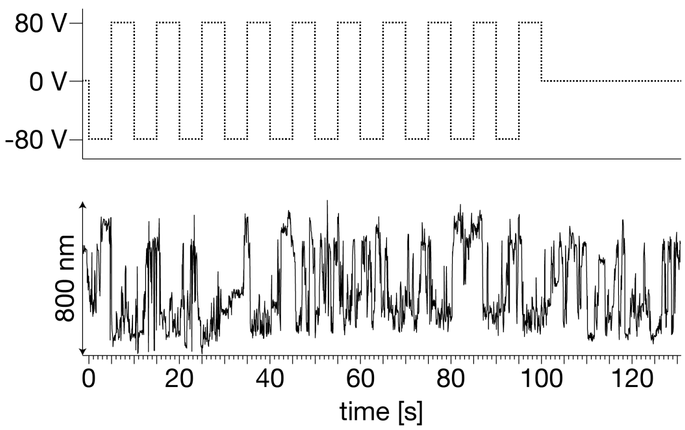


**Figure S 38** Single particle position-time trace of a piston inside of a tunnel which was oriented perpendicularly to an electric-field.

### Supplementary note 1

#### 1.1. Langevin Simulations

The starting point of the theoretical description is the 1D Langevin equation for the diffusion of a particle in an external potential U(x), which reads

$$m\ddot{x}(t) = -\gamma\dot{x}(t) - \nabla U[x(t)] + F_R(t), \qquad \text{(S1)}$$

where m is the effective mass, $\gamma$ is the friction coefficient, -$\nabla U[x(t)]$ is the force acting on the particle and $F_R(t)$ the random force which introduces the stochastic forces from the environment. We assume a stationary equilibrium Gaussian process with white noise, i.e. $< F_R(t) > = 0$ and $< F_R(t)F_R(0) > = B^2\,\delta(t)$, where $B = \sqrt{2k_B\,T\,\gamma}$ .

A periodic sine or cosine potential is the simplest example of a periodic potential. However, the analysis of experimental data in fig. 4C suggests that the effective potential is much sharper than a sine or cosine potential. Hence, we choose a more general periodic potential landscape U of the form

$$U[x(t)] = U_B\left(1 - \sum_{n=0}^{N} e^{-\frac{(x(t)-\tilde{n}a)^2}{2b^2}}\right), \qquad \text{(S2)}$$

where a is the period, b is the width of the Gaussian potential well, $U_B$ is the barrier height and $\tilde{n} = \frac{1+2n}{2}, n \in \mathbb{Z}$. In fig. S1, we show the potential for $U_B = 5\,k_BT$, b = 4 nm and a = 64 nm.

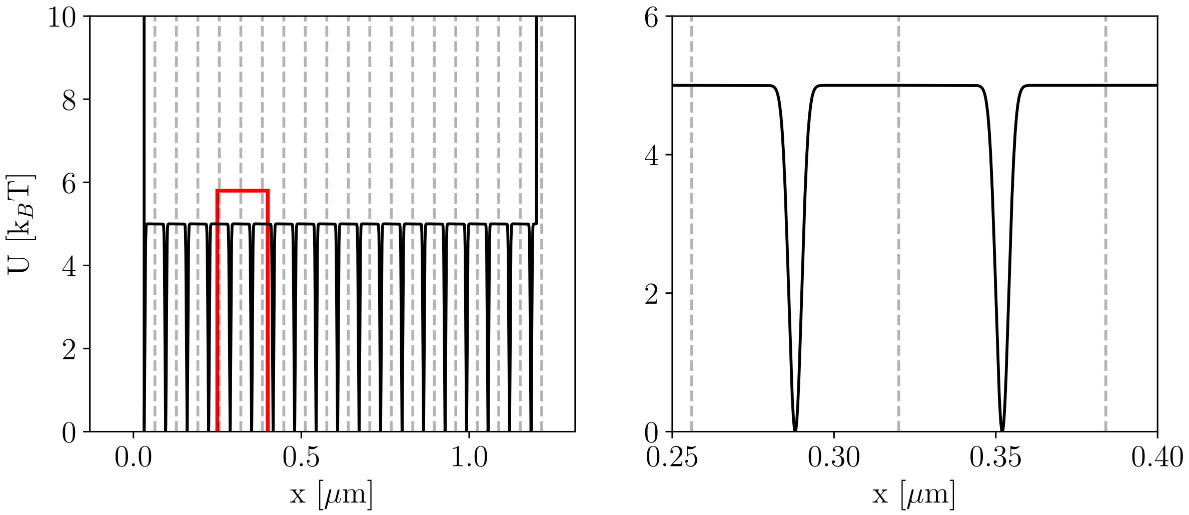


**Figure S 39** Left: Landscape of the periodic Gaussian potential in Eq. S2 with a period of a = 64 nm (see vertical dashed lines), a width of b = 4 nm and a barrier height of $U_B = 5\,k_BT$. Right: Close view of the potential in the area of the red inset in left.

We assume an overdamped motion, so that the Langevin equation in (S1) simplifies to

$$v(t) \equiv \dot{x}(t) = -\frac{1}{\gamma}\nabla U[x(t)] + \frac{1}{\gamma}F_R(t), \quad \text{(S3)}$$

The friction coefficient $\gamma$ for the free diffusion case is estimated from the standard deviation of the averaged experimental velocity distribution in fig. 5E, i.e. $\sigma_{v,exp} \approx 0.59\,\mu m/s$. With $k_B T \approx 4.11 \cdot 10^{-21}$ J at 298 K we obtain:

$$\gamma = \frac{k_B T}{D} = \frac{2k_B T}{\sigma^2_{v,exp}\Delta} \approx 1.39 \cdot 10^5 \,\frac{u}{fs}, \quad \text{(S4)}$$

where $\Delta = 0.1s$ is the experimental time step (frame rate of the detector). Using Einstein's relation for free diffusion, i.e. $\gamma = \frac{k_B T}{D}$, we have a diffusion coefficient of $D \approx 1.77 \cdot 10^{-2}\,\mu m^2/s$. With the experimentally found friction coefficient, we can estimate the inertial time scale $\tau_m = m/\gamma$. With m = 2 MDa, we have $\tau_m \approx 14$ fs for room temperature. This small value suggests that our overdamped assumption in Eq. S3 is valid.

We simulate trajectories by numerically solving Eq. S3 using the fourth-order Runge-Kutta method with a time step $\delta t$. We adjust the potential $U[x(t)]$ to experimental conditions. At positions x = 0 $\mu m$ and x = 1.2 $\mu m$, we add infinitely high potential barriers to the periodic potential landscape (see fig. S 39), which approximately correspond to the length of a filament that is closed at both ends. These boundaries are reflective, which is a realistic assumption for the movement of a piston in the filament. We sample time steps for the random force $F_R$ by computing Gaussian white noise with zero mean and variance

$$\sigma_F^2 = \langle F_R(0)F_R(0) \rangle, \quad \text{(S5)}$$

$$= B^2/\delta t, \quad \text{(S6)}$$

$$= 2k_B\, T\, \gamma/\delta t. \quad \text{(S7)}$$

The mean and variance expression correspond to the choice of random force in the Langevin equation in Eq. S1. At each numerical time step $F_R^i$, we draw a random number from a Gaussian distribution $\mathcal{N}$ with zero mean and a standard deviation of $\sigma_F$ in Eq. S7, i.e. $F_R^i \sim \mathcal{N}(0, \sigma_F)$. We simulate trajectories with a time resolution of $\delta t$ =10 $\mu s$ and a total length of 600 s. The simulated trajectories $x_i$ are discretized and used to calculate the discrete velocity

$$v_i(t) = \frac{x_i(t+\Delta/2) - x_i(t-\Delta/2)}{\Delta}, \quad \text{(S8)}$$

with $\Delta = 0.1\,s$ being the experimental time resolution. In fig. S 40, we show exemplary simulated, discretized trajectories with the potential landscape in Eq. S2 for different barrier heights. We observe that the standard deviation of the velocity probability distribution depends on the barrier height choice. This suggests that the non-Gaussian behavior found in the experimental data arises from the summation of piston traces in periodic potentials with different barrier heights.

For the simulation results we show in fig. 5E, we simulated 100 of these trajectories with randomly chosen potential barrier heights from a uniform distribution in the range between $U_B = 1\,k_B\,T$ and $U_B = 10\,k_B\,T$, a potential period a = 64 nm and a barrier width of b = 4 nm. We kept the potential landscape homogeneous for each simulation, meaning constant barrier height over time and space. Afterwards, we computed the discrete velocity distributions with Eq. S8 to obtain the result in fig. 5E.

The choice of the individual barrier heights was based on estimates from the experimentally found potential landscapes. However, these profiles suffer from limited sampling, and no exact combination of heights could be determined, so random heights from a uniform distribution were chosen. Therefore, we assume that a specific "optimal" combination of barrier heights can completely reproduce the experimental distribution.

##### 1.2. Adding Localization Noise to the Simulations

The simulations based on the Langevin miss an important experimental ingredient. The detection method based on fluorescence videos using super-resolution centroid tracking is subject to a limited resolution. We tracked pistons before adding invader strands. These pistons should be stationary and provide a good basis for estimating the spatial resolution. The spatial resolution can be considered as natural localization noise added to the actual piston position. Regarding the simulation setup, such localization noise of the piston's position can be modeled by adding a Gaussian random variable $x_{loc}(t)$ with zero mean and adjustable standard deviation to the discrete position (*2*). The model prediction for the experimentally measured position reads as $x_{exp}(i\Delta) = x_{sim}(i\Delta) + x_{loc}(i\Delta)$. For the simulation results we show in fig. 5D and fig. S 39, we added localization noise $x_{loc}$ sampled from a Gaussian distribution with zero mean and a standard deviation of $\sigma_{loc} = 5$ nm, i.e. $x_{loc}(i\Delta) \sim \mathcal{N}(0, \sigma_{loc})$, to the discretized trajectories.

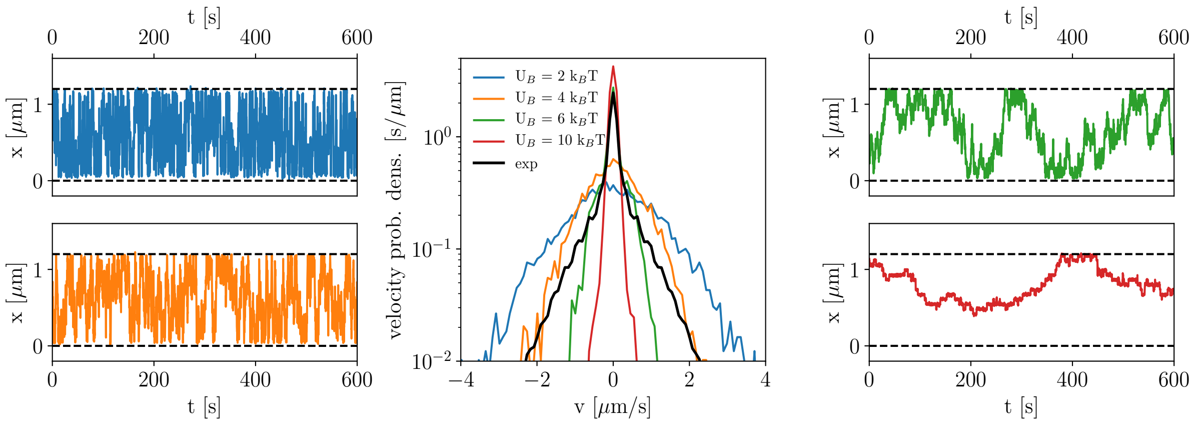


**Figure S40 Examples of overdamped Langevin simulations in a periodic Gaussian potential** (see Eq. S2) with simulation resolution $\delta t$ =10 $\mu s$ and discretized to a time resolution of $\Delta = 0.1$ s, with a period of a = 64 nm, a width of b = 4 nm and varying barrier height $U_B$ (denoted by different colors). For each simulation, the potential landscape was kept homogeneous, meaning with constant barrier height over time and space. Localization noise $x_{loc}(i\Delta)$ sampled from a Gaussian distribution with zero mean and a standard deviation of $\sigma_{loc}$= 5 nm was added to the discretized trajectory positions. Here we used $k_B T$ for 298 K and $\gamma = 1.39 \cdot 10^5 \frac{u}{fs}$ (estimated by Eq. S4). Middle: Calculated discrete velocity probability distributions of the individual simulations (colored, Eq. S8) in comparison with the averaged velocity distribution of the experimental trajectories (black).

### MATERIALS & METHODS

**Design of scaffolded DNA origami objects**

All objects were designed using caDNAno v0.2 (*1*).

**Folding of DNA origami nanostructures**

The folding reaction mixtures for the piston and the two capping objects contained scaffold DNA (*3*) at a final concentration of 50 nM and oligonucleotide strands (IDT Integrated DNA Technologies) at 200 nM each. The folding reaction mixtures for the barrel contained both scaffolds DNA (*3*), (*4*) at a final concentration of 20 nM each and oligonucleotide strands (IDT Integrated DNA Technologies) at 150 nM each. The folding reaction buffers contained 5 mM TRIS, 1 mM EDTA, 5 mM NaCl (pH 8) and 15 mM $MgCl_2$ for the barrel and 20 mM $MgCl_2$ for the piston and caps. The folding reaction mixtures were subjected to different thermal annealing ramps using TETRAD (MJ Research, now Biorad) thermal cycling devices. **Barrel:** 15 minutes at 65°C, followed by three-hour intervals for each temperature, starting at 56°C down to 53°C, decreasing by 1°C every step. **Piston:** 15 minutes at 65°C, followed by one-hour intervals for each temperature, starting at 64°C down to 47°C, decreasing by 1°C every step. **Caps:** 15 minutes at 65°C, followed by one-hour intervals for each temperature, starting at 52°C down to 40°C, decreasing by 1°C every step. Finally, all folding reaction mixtures were incubated at 20°C before further sample preparation steps.

**Purification of DNA origami nanostructures by gel extraction**

All folded objects were purified from excess oligonucleotides by gel-electrophoretic separation (*5*). The samples were electrophoresed on 1.5-2.5 % agarose gels containing 0.5x tris-borate-EDTA and 5.5 mM $MgCl_2$ for 1-3 hours at 70 or 90 V bias voltage in a water-cooled gel box. The desired bands where then cut out of the gel with an X-tracta Generation 2 hand punch. The extracted sample was then centrifuged at 2000 rcf for 5 min in a Freeze ‘N Squeeze DNA Gel Extraction Spin Colum, pore size 0.45 µm (BioRad).

**Buffer exchange of DNA origami nanostructure samples by ultrafiltration**

Buffer exchange after gel extraction was performed via ultrafiltration (Amicon Ultra 0.5 ml Ultracel filters, 50K and 100K) with buffer containing 5 mM TRIS, 1 mM EDTA and 500 mM NaCl (*6*). All centrifugation steps were performed at 10k G for 3-10 minutes at 25°C. The filters were first filled up with 0.5 ml of buffer and centrifuged. 0.5 ml of origami sample were then added and centrifuged. Another 3 rounds of adding 0.45 ml buffer and subsequent centrifugation were performed before a final retrieving step, where the filter inset was turned upside down, placed into a new tube and centrifuged.

**Negative-stain TEM**

5 µl of sample (10-50 nM DNA object concentrations, 20-40 mM $MgCl_2$ concentration and 1-4.5 M NaCl) were pipetted onto a plasma-treated (45 seconds, 35 mA) formvar-supported carbon-coated Cu400 grid (Electron Microscopy Sciences). The sample droplets were incubated for 30 s-10 min on the grids and then blotted away with filter paper. A 5 µl droplet of 2% aqueous uranyl formate (UFO) solution containing 25 mM sodium hydroxide was added and blotted away as a washing step. A 20 µl UFO droplet was then added, incubated for 30-40 s and blotted away. The grids were then air dried for 10-20 minutes before imaging in a Philips CM100, an FEI Tecnai 120 and a Jeol JEM3200 FSC microscope. Automated particle picking was performed with cryYOLO (*7*), 2D class averaging was performed with Relion (*8*).

**JEOL JEM3200 FSC image acquisition**

The JEOL JEM3200 FSC was operated at 300kV and the energy filter slit was set to 20eV width. Images were acquired with an DE64 direct electron detector operated in integration mode at a magnified pixel size of 16A. 75 frames were saved per image and subsequently corrected for residual motion with 10x10 patches using the RELION 3.0.8 implementation of Motioncor2 (*9*).

**Gel electrophoresis**

All samples were electrophoresed on 1.5-2.5% agarose gels containing 0.5x tris-borate-EDTA and 5-25 mM $MgCl_2$ for 1-3 hours at 70 or 90 V bias voltage in a water of ice-water cooled gel box. The loaded samples contained final monomer concentrations of 5-20 nM (unless otherwise noted). The gels were stained with ethidium bromide, if the samples did not include fluorescent dyes and were scanned with a Typhoon FLA 9500 laser scanner (GE Healthcare) at a resolution of 25 or 50 µm/pixel. Resulting images were further processed using Photoshop CS5.

**Transport system sample preparation protocol for fluorescence measurements**

1. Folding of all four monomers at their respective optimal folding conditions, according to initial folding screens.
2. Purification of excess oligonucleotides by gel extraction for all four samples.
3. Addition of a Biotin- modified oligonucleotide to the purified barrel sample to bind to the respective anchor sequence on the outside of the barrel.
4. Four rounds of ultrafiltration with buffer containing 0.5 M NaCl and no $MgCl_2$ for all four samples.
5. Mix the barrel and the piston sample at a barrel to piston ratio of 6:1 – 10:1, incubate at 30°C in 3 M NaCl for 12-16 hours to build dimers.
6. Add oligonucleotide sequences to permanently close the barrel and incubate at 30°C for 15 min.

7. Incubate the closed barrel-piston dimers at 40°C in 2 M NaCl for 1 hour to reduce possible aggregation due to high salt conditions.
8. Add the first set of oligonucleotide sequences to trigger the polymerization of piston-loaded and empty barrels at 4:1 staple to barrel ratio. Incubate at 40°C for 30-60 min.
9. Add the second set of oligonucleotide sequences for the polymerization reaction at 4:1 staple to barrel ratio. Incubate at 40°C for 30-60 min.
10. Add the first cap at a cap to tunnel ratio of 6:1, assuming that the average tunnel consists of roughly ~30-40 barrels. Incubate at 40°C in 2 M NaCl for 1 hour.
11. Add the second cap at the same cap to tunnel ratio and incubate at 40°C in 2 M NaCl for 1 hour.

The entire apparatus construction protocol is complete within 20 hours, starting with purified monomers (peg, barrel and both caps) and finishing with fully functional transport systems, ready for real-time fluorescence experiments.

**Single-molecule fluorescence microscopy experiments for free diffusion**

1. Cover slides (Sigma Aldrich) were cleaned in 2 M NaOH for 30 min and then sonicated in 2% Hellmanex, rinsed with double distilled water ($ddH_2O$), then sonicated in $ddH_2O$, again rinsed with $ddH_2O$ and finally sonicated in ethanol (99%). All sonication steps were performed for 5 min. The slides were then dried at 70°C for 1 hour. A solution of 0.5% bioPEG-silane (solved in ethanol) with 1% acetic acid was incubated on the slides at 70°C for 30 min. The slides were then rinsed with $ddH_2O$ and flushed with $N_2$ until dry and stored protected from light (*10*).
2. Place a reaction chamber on top of the cover slides and wash with NeutrAvidin (0.05 mg/ml) in T50 buffer (10 mM Tris, 50 mM NaCl). Incubate for 10-15 min and wash with buffer containing 2 M NaCl.
3. Add the sample containing capped tunnels with pistons trapped inside to the reaction chamber and bind the tunnels to the pegylated glass surface via biotin-neutravidin interaction. Incubate for 10-30 min depending on the desired surface coverage.
4. Wash the reaction chamber with buffer containing an oxygen scavenging system (100 mM tris-HCl, 2 mM Trolox, 0.8% D-glucose, catalase (2000 U/ml), glucose oxydase (165 U/ml), purchased from Sigma Aldrich) and 1 M NaCl, along with invader oligonucleotide sequences at final concentrations of 1 μM to release the pistons from their initial docking sites. Incubate at room temperature for 15 min.
5. Acquire movies at 20-35 °C in a custom-built objective-type TIRFM (oil-immersion objective, 100x, apochromat, NA 1.49; Nixon with an acousto-optical tunable filter (Pistonasus Optics)). Green (532 nm, Oxxius) and red (640 nm; Oxxius) diode lasers were used to excite the Cyanine-3 and Cyanine-5 dyes attached to the pistons and barrels. Their fluorescence signals were divided with dichroic mirrors and detected by two EMCCD cameras (Andor iXon+). Movies were acquired in 512x512 pixel format with a frame rate of 10 fps per channel. A piezo-

driven sample stage (PInanoXYZ; Physik Instrumente) was used to manoeuvre to different positions on the cover slide. A custom written LabView routine was used to control all microscope components (*11*).

**Single particle centroid tracking**

The piston diffusion was tracked using the ImageJ particle tracker plugin (*12*).

**Temperature control**

Temperature controlled single molecule TIRF measurements were performed using VAHEAT-Micro heating system (Interherence GmbH).

**Single-molecule fluorescence microscopy setup for driven diffusion**

For measurements with applied electric fields, a separate single molecule TIRFM setup was used. The setup was custom-built on the basis an Olympus IX71 microscope body. Three laser light sources with wavelengths 642 nm (Toptica iBeam smart, diode laser, 150 mW, Gräfelfing, Germany), 532 nm (Oxxius 532-50, diode-pumped solid-state laser, 50 mW, Lannion, France), and 488 nm (Toptica iPulse, diode laser, 20 mW) are used as excitation light source. The 488 nm excitation was not used for experiments in this project.  All imaging is performed with a 100x oil immersion objective (UAPON 100xOTIRF objective, NA 1.49 oil, Olympus, Japan). The sample is supported by a piezo z-stage (Physik Instrumente (PI) GmbH, Karlsruhe, Germany). The filter cube was configured with a ZT532/640RPC dichroic mirror and a ZET532/640 (Chroma Technology, Olching, Germany) emission filter. Videos were acquired with an ORCA-Fusion Digital CMOS camera (Hamamatsu Photonics, Japan) attached to the left camera port.

The flow chamber, in which the structures were exposed to the applied fields is assembled from three parts (fig. S41). A top part made from $Al_2O_3$, double-sided adhesive tape 3M 467MP (3M Company, Maplewood, Minnesota, USA) and cover slip. The top part creates buffer reservoirs and helps to attach a custom-made plug. The plug secures 0.2 mm thick platinum wires, to which the operating voltage is applied. The adhesive tape is cut with a laser engraver (Trotec Speedy 100, Trotec Laser, Marchtrenk, Austria) to create a 50µm high channel connecting the reservoirs when sandwiched between top part and cover slip. The applied voltage was controlled by a custom-build LabView routine that supplied control voltages to a custom-built operational amplifier to generate the final output voltage.

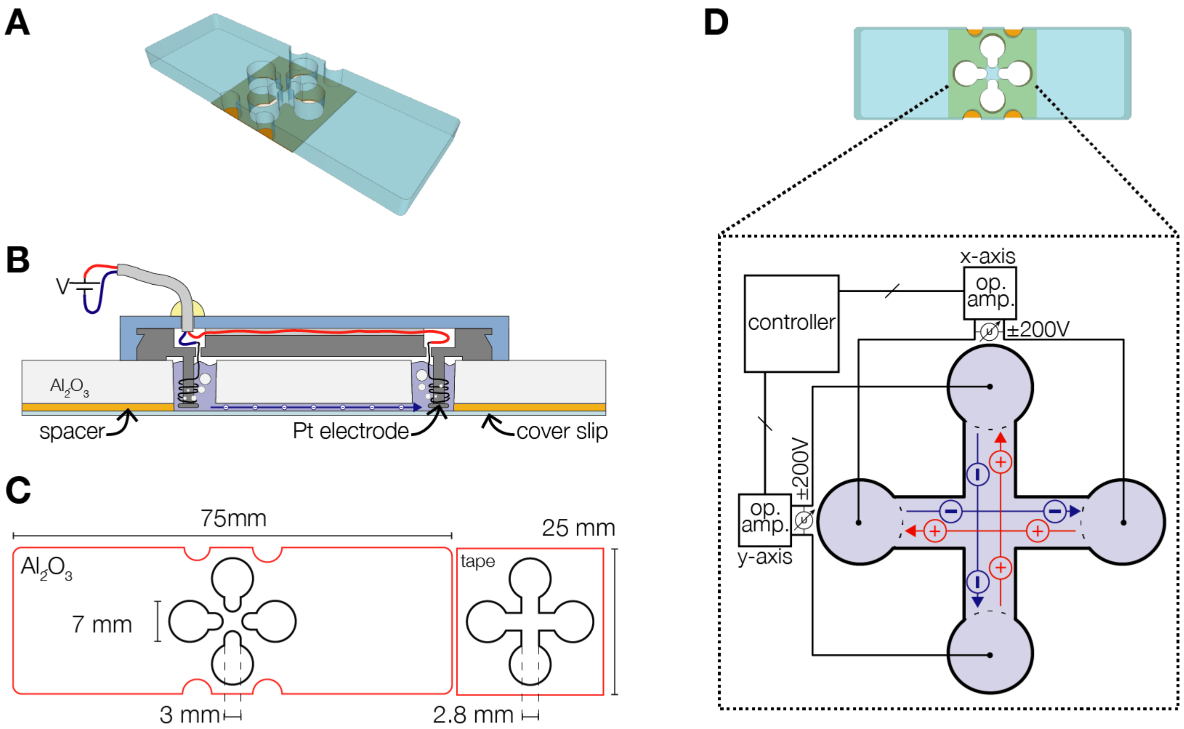


**Figure S41 Electrode setup for the application of electric fields.** (A) Illustration of assembled sample chamber in isometric view. (B) Cross section of sample chamber with inserted plug. For better visibility, material thickness not drawn to scale. (C) Dimensions of top component of sample chamber and adhesive tape. (D) Illustration of assembled sample chamber in top view and schematic of the resulting channel shape and electrode wiring.

#### Captions for movies S1 to S7

**Movie 1**

Exemplary fluorescence microscopy movie with released pistons (merged channels). Real-time. 50 x 50 µm.

**Movie 2**

Exemplary fluorescence microscopy movie with released pistons (merged channels). Real-time. Each pixel is 104 nm.

**Movie 3**

Exemplary fluorescence microscopy movie with released piston (merged channels). Real-time. Each pixel is 104 nm.

**Movie 4**

Longest range: Exemplary fluorescence microscopy movie with released piston (merged channels). Real-time. Each pixel is 104 nm.

**Movie 5**

Fastest particle: Exemplary fluorescence microscopy movie with released piston (merged channels). Real-time. Each pixel is 104 nm.

**Movie 6**

Electric-field driven motion: Exemplary fluorescence microscopy movie with released piston (merged channels). Real-time. Each pixel is 130 nm.

**Movie 7**

Electric-field driven motion: Exemplary fluorescence microscopy movie with released pistons (merged channels). Real-time. Each pixel is 130 nm.
